# Meta-imputation of transcriptome from genotypes across multiple datasets using summary-level data

**DOI:** 10.1101/2021.05.04.442575

**Authors:** Andrew Liu, Hyun Min Kang

**Affiliations:** Department of Biostatistics and Center for Statistical Genetics, University of Michigan, Ann Arbor, Michigan, United States of America

**Author notes:** Contact (AL), (HK).

## Abstract

Transcriptome wide association studies (TWAS) can be used as a powerful method to identify and interpret the underlying biological mechanisms behind GWAS by mapping gene expression levels with phenotypes. In TWAS, gene expression is often imputed from individual-level genotypes of regulatory variants identified from external resources, such as Genotype-Tissue Expression (GTEx) Project. In this setting, a straightforward approach to impute expression levels of a specific tissue is to use the model trained from the same tissue type. When multiple tissues are available for the same subjects, it has been demonstrated that training imputation models from multiple tissue types improves the accuracy because of shared eQTLs between the tissues and increase in effective sample size. However, existing joint-tissue methods require access of genotype and expression data across all tissues. Moreover, they cannot leverage the abundance of various expression datasets across various tissues for non-overlapping individuals.

Here, we explore the optimal way to combine imputed levels across training models from multiple tissues and datasets in a flexible manner using summary-level data. Our proposed method (SWAM) combines arbitrary number of transcriptome imputation models to linearly optimize the imputation accuracy given a target tissue. By integrating models across tissues and/or individuals, SWAM can improve the accuracy of transcriptome imputation or to improve power to TWAS without having to access each individual-level dataset. To evaluate the accuracy of SWAM, we combined 49 tissue-specific gene expression imputation models from the GTEx Project as well as from a large eQTL study of Depression Susceptibility Genes and Networks (DGN) Project and tested imputation accuracy in GEUVADIS lymphoblast cell lines samples. We also extend our meta-imputation method to meta-TWAS to leverage multiple tissues in TWAS analysis with summary-level statistics. Our results capitalize on the importance of integrating multiple tissues to unravel regulatory impacts of genetic variants on complex traits.

**Author Summary:** The gene expression levels within a cell are affected by various factors, including DNA variation, cell type, cellular microenvironment, disease status, and other environmental factors surrounding the individual. The genetic component of gene expression is known to explain a substantial fraction of transcriptional variation among individuals and can be imputed from genotypes in a tissue-specific manner, by training from population-scale transcriptomic profiles designed to identify expression quantitative loci (eQTLs). Imputing gene expression levels is shown to help understand the genetic basis of human disease through Transcriptome-wide association analysis (TWAS) and Mendelian Randomization (MR).

However, it has been unclear how to integrate multiple imputation models trained from individual datasets to maximize their accuracy without having to access individual genotypes and expression levels that are often protected for privacy concerns. We developed *SWAM* (Smartly Weighted Averaging across Multiple datasets), a *meta-imputation* framework which can accurately impute gene expression levels from genotypes by integrating multiple imputation models without requiring individual-level data. Our method examines the similarity or differences between resources and borrowing information most relevant to the tissue of interest. We demonstrate that SWAM outperforms existing single-tissue and multi-tissue imputation models and continue to increase accuracy when integrating additional imputation models.

## Introduction

Genome wide association studies (GWAS) have been able to identify numerous associations between genetic variants and complex traits. However, interpreting the biological mechanisms underlying the association signals remains a challenge [1]. Recently, studies involving gene expression have become increasingly popular as a means to provide biologically meaningful insight into statistical associations [2,3]. Transcriptome-wide association studies (TWAS) is a widely used method to translate GWAS association signals into more interpretable units by examining the association between phenotypes and gene expression levels imputed from genotypes. Associations identified from TWAS can be interpreted as potentially causal relationships between the traits and the genes through gene regulation [4–6]. While TWAS may not detect associations driven by functional mechanisms irrelevant to gene regulation, it increases the specificity and interpretability in identifying GWAS signals driven by gene regulation. Imputed gene expression can be utilized in various contexts of association analysis beyond TWAS, such as Mendelian randomization [7,8] or estimation of trait heritability attributable to cis-eQTLs [9]. Since genotype data from DNA is far easier and cheaper to obtain than expression data from tissues, TWAS based on imputed expression offers excellent augmentation to study the genetic component of gene regulation in addition to RNA-seq-based studies.

The first-generation methods to impute gene expression levels from genotypes train the model from a single-tissue dataset comprising of many individuals with both genotypes and expression profiles [2,3]. For example, a widely-used method PrediXcan [2] uses Elastic net regularization to identify cis-eQTLs (expression quantitative loci) to train the model to impute gene expressions from genotypes. Other methods, such as TWAS [3], employ different regularization but typically produces a linear model to impute gene expressions as a weighted sum of cis-eQTL genotypes. Imputation models are trained using these methods from various population-scale transcriptomic datasets, such as the Genotype-Tissue Expression (GTEx) project [9,10], Depression Genes and Network (DGN) study [11], and The Cancer Genome Atlas (TCGA) [12], and these models are made available in public repository such as predictDB (http://predictdb.org/) or FUSION (http://gusevlab.org/projects/fusion/) so that expression imputation or TWAS can be performed from any genotyped individuals.

Although these first-generation methods for transcriptome imputation have been quite useful, they have limited accuracy mostly due to limited sample size in the training datasets where both genome-wide genotypes and transcriptome-wide expression levels are available. While millions of individuals have been genotyped or sequenced to date [13–16], the sample-size of current population-scale transcriptome data are typically limited only to hundreds or thousands [17] (with the largest study cohort having around 30k participants [18]), primarily due to the difficulty in collecting high quality tissues (other than whole blood) from living donors. Moreover, transcriptomic datasets are prone to potential batch effects between studies [19–22], making it difficult to integrate across multiple datasets to build a large and harmonized resource to be trained from. Furthermore, there are hundreds or thousands of different types of tissues or cells, requiring orders of magnitude larger effort to comprehensively profiles transcriptomes in population-scale across tissues, as in GTEx Project.

Recently, methods to address the shortcomings of the first-generation methods have been developed. When transcriptomic profiles are available across many tissues, such as in GTEx Project, transcriptome imputation can improve by leveraging the shared genetic components across tissues. Even though each tissue represents a unique transcriptomic profile, a large fraction of eQTLs are shared across tissues [23], and the availability of multiple expression measurements across tissues can help more precisely identify the shared eQTLs, which in turn can improve the imputation accuracy. For example, UTMOST trains a transcriptome imputation model a simultaneously across all tissues using a combination of L1 and L2 penalization across markers and tissues, respectively [24]. Another multi-tissue approach, MultiXcan, does not impute transcriptomes, but performs a multi-tissue TWAS across all tissues by including each tissue-specific imputed expression as a predictor variable to improve power to identify association between a trait and a gene, in which the underlying mechanism potentially involves multiple tissues or cell types [25].

Even though UTMOST substantially improves the accuracy of transcriptome imputation, it assumes that expression measurements across multiple tissues are available for overlapping set of genotypes individuals for training imputation models. While this assumption can be met when training from the GTEx dataset (assuming granted access to the individual-level data), it may not be realistic in other circumstances where expression measurements are available for non-overlapping individuals (such as in TCGA), or it is infeasible to obtain individual-level genotypes and expression data due to limited access privilege. As population-scale transcriptomic resources are rapidly increasing, it should be possible in principle to integrate these resources to better impute transcriptomes. While there have been additional methods which have been developed to increase the accuracy of gene expression or TWAS [25–28], none of them – to the best of our knowledge – are able to perform “meta-imputation”, which systematically integrates multiple imputation models without the need to access to individual-level data.

Here we propose Smartly Weighted Averaging across Multiple datasets (SWAM), a multi-tissue transcriptome imputation method based on a flexible meta-analysis across multiple imputation models. Unlike UTMOST, SWAM does not require access to all genotypes and expression datasets for training its imputation model. Instead, it takes individual transcriptome imputation models trained from individual tissues while optimizing the expected imputation accuracy for a target tissue. Moreover, it can seamlessly integrate imputation models trained from multiple datasets comprising of different individuals and tissues. As a result, SWAM can integrate across hundreds of imputation models across GTEx, DGN, and TCGA projects without requiring all individual-level data to substantially improve the imputation accuracy over existing methods, as we demonstrate with GEUVADIS data. Moreover, we demonstrate that SWAM improves the power of TWAS over single-tissue methods and many alternative multi-tissue methods.

## Results

### Smartly Weighted Averaging across Multiple Datasets (SWAM)

We propose *<u>S</u>martly <u>W</u>eighted <u>A</u>veraging across <u>M</u>ultiple datasets* (SWAM), a method that provides a flexible framework to impute tissue-specific expression by integrating single-tissue imputation models across many tissues and datasets (**Figure 5**). The key principle behind SWAM is to improve the accuracy of transcriptomic imputation by determining the optimal linear combination of multiple imputation models in terms of expected imputation accuracy. To do this, SWAM requires a reference tissue (tissue of interest) to be defined as a basis to determine the relative contributions from multiple imputation models. Using the individual-level genotypes and expression of only the reference tissue, SWAM integrates imputation models trained from different tissues and datasets (e.g. GTEx, DGN, and TCGA) without requiring individual-level data except for the reference tissue.

**Figure 5.**
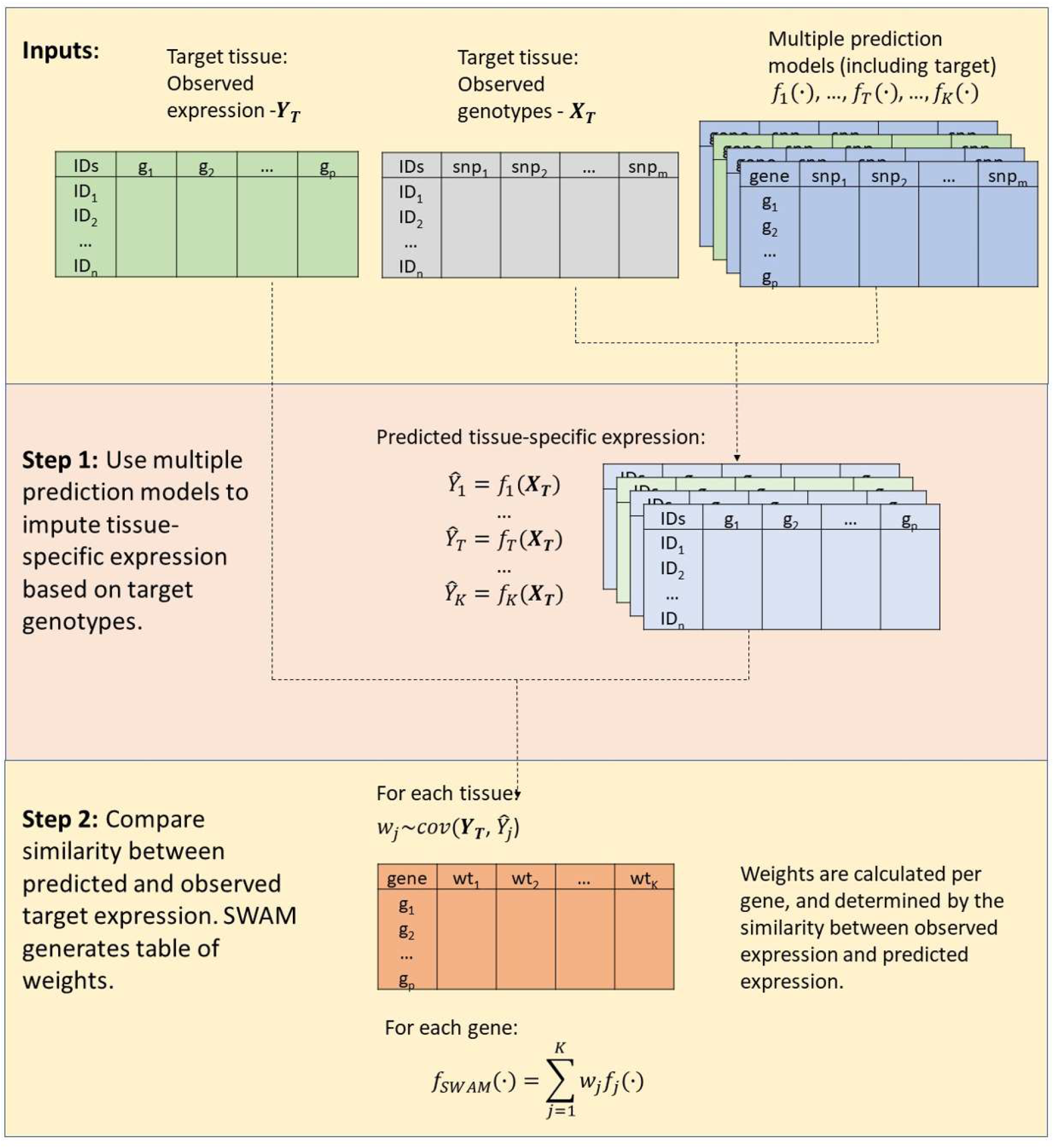
overview of SWAM method. This figure demonstrates the training of the imputation model using the reference data. The inputs required for SWAM are a set of reference genotypes with sample matched measured expression, and the multiple imputation models to be included. The list of multiple imputation models must also include a model derived from the reference data, which can be done via prediXcan. SWAM uses these models to impute tissue-specific expression levels from the reference genotypes. These imputed expression sets are then compared with the measured expression of the reference set. The weights are calculated based on the similarity between the measured and imputed expression and the covariance structure of tissues. For full details, see the methods section.

The first step of SWAM is to apply each transcriptomic imputation model to the reference genotypes, which results in individual-level, tissue-specific imputed expression. The second step of SWAM compares each imputed expression with the measured expression of the reference tissue to calculate optimal weights by linearly combining multiple imputation models to maximize expected mean squared error (MSE) (see Methods for the details). The output of second step is an integrated transcriptomic imputation model compatible with the PrediXcan and MetaXcan software tools. Using this SWAM model, we can impute the transcriptome of any samples of interest with genotype information available (via PrediXcan), or to use the model and covariance matrix directly to perform TWAS (via MetaXcan) when GWAS summary statistics are available (S1 Fig).

### Simulation study demonstrates the robustness of SWAM across various scenarios

We performed simulation studies to evaluate SWAM’s ability to robustly impute expression by leveraging tissue-specific and cross-tissue components across a wide spectrum of parameter settings. To do this, we independently simulated multi-tissue expression levels along with genotype data for both our training and validation sets (see Materials and Methods). We compared SWAM with two heuristic approaches – *naïve average,* which equally weights individual tissue and *best tissue,* which only uses the tissue with the highest expected imputation accuracy – as well as with *single-tissue* imputation.

As expected, we observed *naïve average* to be particularly powerful when the causal variants are shared across all relevant tissues (**Figure 6A**), identifying 94% of genes as significantly imputable at FDR < 0.05. When all causal variants were tissue-specific, the naïve average only identified 25% of genes to be imputable. On the other hand, best-tissue was more powerful (38%) than naïve-average when the all causal variants were tissue-specific, but worse when all causal variants are shared. When only *single-tissue* was used for imputation, the performance stayed similar regardless of the tissue-specificity. Encouragingly, SWAM outperformed all three methods across all ranges of tissue-specific and cross-tissue heritability settings. We believe this is because SWAM learns tissue-specific weights without pre-conceptions of tissue relatedness, and thus determines the weights for relevant tissues while ignoring unrelated ones.

**Figure 6.**
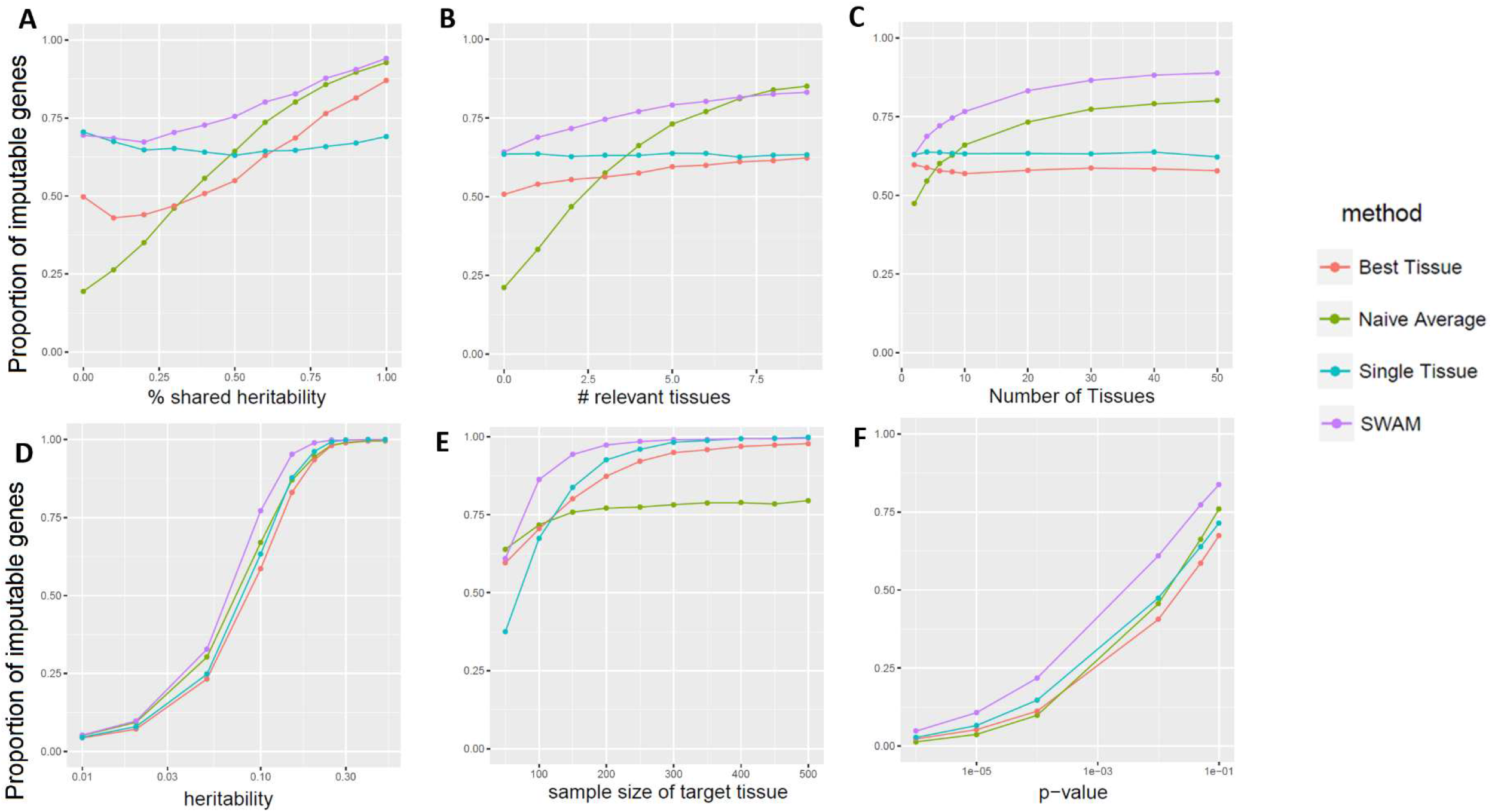
simulation study comparing SWAM with naïve average, best tissue and single tissue methods. We ran each simulation 10,000 times, with the following default settings: 10 total tissues (1 target, 4 relevant, 5 irrelevant),100 SNPs (2 per tissue), 10% genetic heritability, 50% shared heritability between relevant tissues. In addition, the sample size of the target tissue was 100 individuals, and the remaining tissues had 200 individuals. This was done to emphasize the importance of integrating information from other tissues when the quality of the target tissue model is limited. In panel (A),we varied the number of relevant tissues, from 0 to 10. Panel (B)shows the improvement when the total number of tissues is increased, with the number of irrelevant tissues fixed at 50% of the total. Panel (C) showsthe effects of changing the shared heritability for the relevant tissues. We note here, that each tissue has 2 causal SNPS – for the relevant tissues, 1 of these causal SNPS is shared with the target tissue while the other is independent of all simulated tissues. Panel (D)shows the performance of the approaches for different levels of genetic heritability. This simulation demonstrates the range of heritability that we would expect to see the most improvement. Empirically, we do notice the same trend seen here, as SWAM performs similarly the single tissue model when the cross-validated R-squared is high. Panel (E)shows the effects of target tissue sample size. The x-axis pertains to the sample size of the target tissue only, and all other tissues were fixed at 200 individuals. Finally, panel shows the performance of the methods at different p-value thresholds, using the default simulation settings.

A similar trend is observed when we vary the number of relevant tissues that shares cross-tissue heritability (**Figure 6B**). In the case where there are no relevant tissues other than the target tissue, naïve average is least powerful while SWAM performs as well as the *single tissue* approach. This suggests that in this scenario, SWAM is correctly giving non-zero weights to only the target tissue, making it similar to the *single-tissue* method. In the other scenario where every tissue is relevant, SWAM provides a similar power to the *naïve average* approach, suggesting that SWAM is robustly assigning weights to each relevant tissue. Similarly, when there are more tissues available in overall (assuming 50% are relevant tissue sharing cross-tissue heritability), the power of SWAM and *naïve average* keep increasing while *single-tissue* and *best-tissue* remain similar (**Figure 6C**).

Our simulation study also evaluated the impact of sample size of the reference tissue. We hypothesized that *single-tissue* would perform poorly when the sample size of the reference tissue was small, which was indeed observed in our results (**Figure 6D**). When the reference tissue has sample sizes of 50, 100, 200, we observed that *single tissue* method identified 36%, 66%, and 92% of imputable genes. Because additional tissues are helpful especially when the reference tissue has smaller sample size, the *best tissue* approach performed better at lower sample size (59% at n=50), but worse at higher sample size (88% at n=200). Similarly, *naïve average* also performed better at lower sample size (63% at n=50), but worse at higher sample size (78% at n=200). However, SWAM consistently outperformed single tissue across all cases (59%, 86%, 97% at n=50, 100, 200). This implies that borrowing information from a relevant tissue (to the reference) is useful in these situations and SWAM robustly estimates the weights from each tissue accounting for the uncertainty from different sample sizes.

### SWAM outperforms other transcriptome imputation methods in evaluations with real data by considering the bias-variance tradeoff

We applied SWAM to create multi-tissue imputation models from GTEx v6, using lymphoblastoid cell lines (LCL; the official tissue name in GTEx was “Cells – EBV-transformed lymphocytes”) as the reference tissue, to evaluate its imputation accuracy of LCL transcriptomes of 344 European samples from the GEUVADIS consortium [29]. We compared the accuracy of SWAM with various methods, including *single tissue* imputation models (generated by PrediXcan), *naïve average*, *best tissue*, and another multi-tissue method *UTMOST*.

Among the single-tissue imputation models, we observed that the imputation from LCL identified 1,552 genes as significantly imputable at FDR < 0.05. Interestingly, we observed that another tissue, fibroblast cell lines (FCL; the official tissue name in GTEx was “Cells – Cultured fibroblasts”), identified even more genes (1,690 genes) as significantly imputable for GEUVADIS LCL expression levels. One of the outstanding differences between LCL (n=114) and FCL (n=272) models were the sample size used for training. We suspect that this is due to (1) the difference in sample size (i.e., FCL imputation has less variance) and (2) the similarity of transcriptomic profiles between LCL and FCL (i.e., FCL model tends not to introduce large bias). However, tissues with larger sample size did not always result in more accurate imputation. When we examined the results from Skeletal muscle model (n=361), which had the largest sample size in GTEx v6, we identified only 1,197 genes as significantly imputable. This is likely because the large differences of transcriptomic profiles between LCL and Skeletal muscle (i.e., Skeletal muscle model tends to introduce large bias). These examples demonstrate that both sample size and tissue relevancy are important for maximizing the imputation accuracy. In statistical terms, our primary interest was to reduce the mean-squared error (MSE), which is the sum of Bias^2^ and Variance. We suspect that FCL model performed better than LCL models due to much smaller variance (because of larger sample size), and better than Skeletal muscle models due to much smaller bias (S2 Fig). We hypothesized that by combining imputations from multiple models, we can minimize MSE by substantially reducing variance without introducing excessive bias, which was our main motivation for developing SWAM.

When evaluating the multi-tissue methods, our two heuristic approaches, *best-tissue* and *naïve average* identified 2,493 and 2,666 significantly imputable genes, respectively, which was >47% and >57% higher than any single tissue models. *UTMOST* (using LCL as the reference) also substantially increased the number of imputable genes (2,238 genes, >32% increase over any single tissue), but surprisingly, it had fewer than the imputable genes compared to the two heuristic approaches. Finally, when we applied *SWAM* specifying GTEx LCL as the reference tissue, the number of imputable genes further increased to 3,040, which is >79% larger than any other single tissue models (S1 Table, **Figure 7A**). Interestingly, *SWAM* improved the imputation accuracy over *UTMOST* even though it requires individual-level data only for one tissue (i.e., LCL) in GTEx while *UTMOST* requires simultaneous access to individual-level data across all tissues. These results demonstrate that SWAM offers an accurate and flexible meta-imputation framework by optimally combining multiple imputation models across tissues.

**Figure 7.**
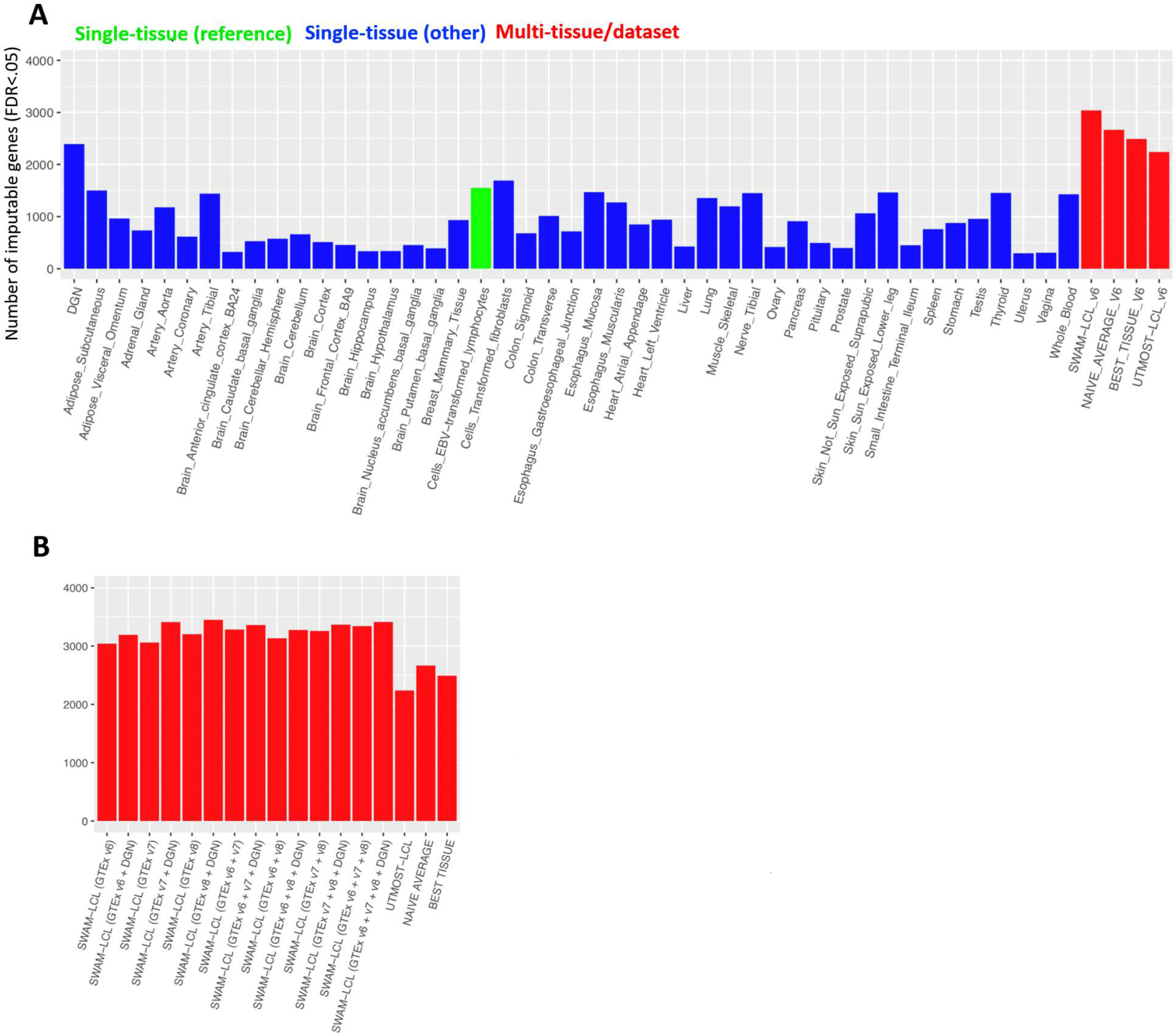
Empirical validation of SWAM using lymphoblastoid-cell line data from GEUVADIS consortium. We used our LCL-targeted SWAM model to impute expression levels based on the genotypes of 344 European samples. We then calculated the concordance between imputed expression and measured LCL expression. We repeated this for all of the other methods mentioned here. (A) shows the performance of SWAM against the single-tissue models from 44 tissue-specific predictDB models derived from GTEx version 6. In (B), we derived various SWAM models using every combination of the following: 1) all GTEx v6 tissues, 2) all GTEx v7 tissues, 3) all GTEx v8 tissues, and 4) Depression Gene Network (DGN) single tissue whole blood model from predictDB. Here, we also included the UTMOST LCL model, naïve average and best tissue models, all derived from GTEx v6.

### SWAM enables meta-imputation of expression levels across multiple heterogeneous datasets

One of the important advantages of SWAM compared to other multi-tissue imputation methods is the ability to integrate imputation models across heterogeneous datasets where samples may not necessarily overlap. To evaluate the benefit of SWAM’s ability for multi-dataset “meta-imputation”, we integrated imputation models trained from GTEx v7 and v8, as well as 922 whole blood transcriptomes from Depression Gene Network (DGN). The rationale to include GTEx v7 and v8 models is that the datasets are slightly different from v6 (for example, v7 has more samples in all tissues except for LCL, FCL, and whole blood) and integrating multiple training models from slightly different versions of datasets may improve the accuracy. The reason to include DGN whole blood is that the sample size is much larger than any individual tissue GTEx, so it may help further reduce the variance and MSE of the imputation model.

When applying SWAM to GTEx v6, v7, or v8 datasets individually, the number of significantly imputed genes at FDR < .05 were 3,040, 3,060, and 3,203, respectively (**Figure 7B**). However, when all datasets were combined, the number of imputable genes increased to 3,342. These results suggest that imputation across multiple datasets can help even when the datasets are highly overlapping. When we additionally integrated SWAM with the DGN whole blood model, which detected 2,390 imputable genes by itself, the number of imputable genes by the integrated SWAM model further increased to 3,413. Note that we needed individual-level data only for the reference tissue/data (GTEx v6 LCL in our experiment), so an arbitrary combination of imputation models, which consist of only summary-level data, can be seamlessly added to the meta-imputation framework of SWAM.

Overall, using all 49 GTEx v8 tissues in combination with the DGN whole blood model provided the highest number of imputable genes, with a 112.9% improvement over the corresponding GTEx v8 PrediXcan-LCL model (single tissue), and a 13.5% improvement over the GTEx v6 version of SWAM-LCL (multi-tissue) (**Figure 7B**). Regardless of the version of GTEx used, including the DGN whole blood model gives a substantial improvement in number of imputable genes compared to not including it in the model. Another interesting observation is that while PrediXcan-LCL (v6) appears to perform better than PrediXcan-LCL (v7), SWAM-LCL derived from v7 performs better than v6 SWAM-LCL. This may suggest that while GTEx v7 PrediXcan-LCL may not have had a significant improvement in eQTL detection compared to its predecessor, other tissues may have improved in more substantial ways. This is because the sample size for LCL in v7 decreased by 18 samples, whereas other non-blood tissues had substantial sample size gains of up to 89 individuals. Here, SWAM leverages the increase in quality from other tissues, which allows for better overall imputation regardless of the quality of the target tissue itself.

### SWAM robustly captures both tissue-specific and cross-tissue regulatory components

The key component behind the robust performance of SWAM is that it learns how to distribute weights across multiple imputation models for each gene individually. If a gene shares eQTLs across many tissues, the SWAM’s weights will be distributed evenly across tissues and the model will behave similarly to the naïve average heuristic. For example, *ERAP2* is a well-known gene with shared eQTLs profiles across most tissues. In the GTEx (v6), *ERAP2* can be reliably imputed with any of the 44 single-tissue imputation models from PrediXcan with r^2^ > 0.77 or more eQTLs. As a result, the weights from SWAM is almost evenly distributed across the tissues, ranging from 0.018 to 0.027 (S2 Fig), and the accuracy of SWAM (r^2^ = 0.795) is very similar to the accuracy of naïve average (r^2^ = 0.796).

On the other hand, when the imputation model from the reference tissue is not particularly good due to smaller sample size or other technical issues, SWAM can substantially improve accuracy by leveraging eQTL sharing from other tissues. For example, the single-tissue imputation accuracy of *GSTM1* is relatively low in LCL tissue (r^2^ = 0.368) compared to the accuracy of the 38 other tissues in which a PrediXcan imputation model is available (average r^2^ = 0.61). Using SWAM, the predictive R-squared increases to r^2^ = 0.741 by assigning positive weights to 31 tissues (S2 Fig).

Finally, for genes that are highly tissue-specific, the SWAM’s weights will be distributed similarly to the best tissue heuristic. For example, CTSK is expressed in most tissues, but has eQTLs in only 16 tissues, (S2 Fig). SWAM assigns weights to 7 of these tissues, and substantially improves the predictive accuracy from r^2^ = 0.111 to r^2^ = 0.447.

### Comparison of imputation models in the context of TWAS

We conducted TWAS analysis using SWAM, UTMOST, and PrediXcan models via MetaXcan [30]. In addition, we also used S-MultiXcan [25] to simultaneously test all of the PrediXcan models using their PCA regression approach. We used a Bonferroni correction to establish p-value threshold for each analysis separately, based on the number of genes imputed. Overall, we found that among the methods that directly estimate expression levels (SWAM, UTMOST, PrediXcan), SWAM outperformed the other methods in terms of number of associations detected (see S4, S5, S6 Tables). For example, PrediXcan models on average detected 23.7, 23.2 and 4.0 transcriptome-trait associations for HDL, LDL and T2D respectively. For SWAM, we observed an average of 79.7, 77.8 and 8.4 associations per tissue, whereas UTMOST yielded an average of 69.3, 61.6 and 8.8 associations per tissue, for the three traits respectively.

We plotted transcriptome-wide signals for the LDL trait using the GTEx v6 liver model for PrediXcan, UTMOST and SWAM (**Figure 8**). One interesting signal gained from the SWAM analysis is the APOC1 gene, which is primarily expressed in the liver and has been implicated in playing a role in HDL and LDL/VLDL (very low-density lipid) metabolism [31].

**Figure 8.**
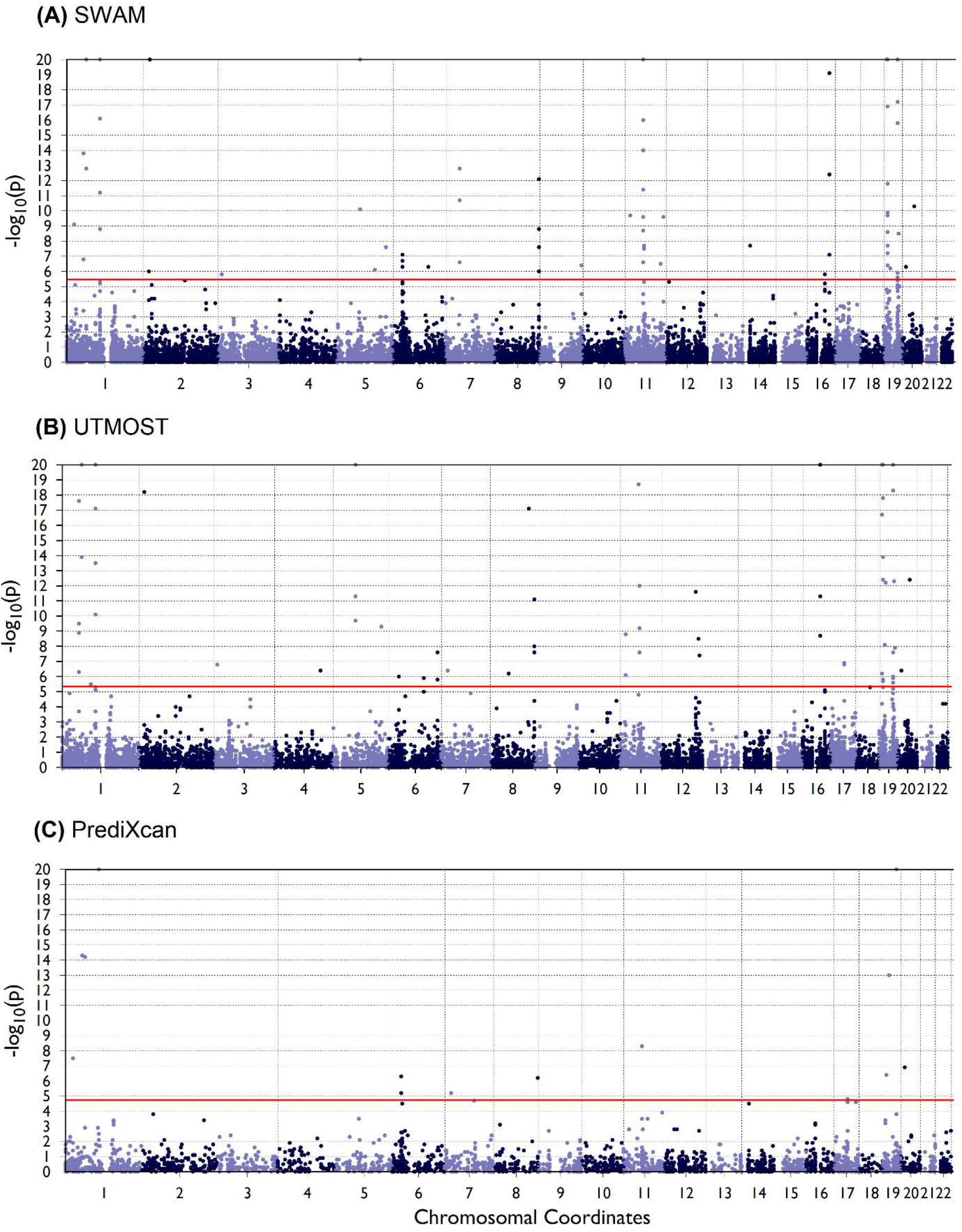
TWAS on LDL trait targeting liver using SWAM, UTMOST, and PrediXcan models. TWAS was performed using metaXcan on the LDL trait from the Global Lipids Genetics Consortium (GLGC) GWA analysis. For a consistent comparison, the SWAM and UTMOST models were derived from GTEx version 6 tissues, and the prediXcan model used was GTEx v6 liver. The number of associations were: 74, 69 and 19 for SWAM, UTMOST and prediXcan respectively. P-values were capped at 10^−20^ in these plots.

One potential shortcoming for both multi-tissue approaches (SWAM and UTMOST) appear to be that the number of unique signals (across all tissues) is fewer than those generated by PrediXcan’s single tissue models. For example, SWAM produces 210 unique associations for the HDL trait, while we see 187 unique associations from UTMOST and 248 unique associations from PrediXcan. Similarly, MultiXcan detects 284 significant associations when scanning across all tissues (based off the PrediXcan models). It appears that while the multi-tissue methods can leverage information from other tissues to impute expression accurately, marginal association signals in TWAS are potentially lost using these approaches. However, we found that a high number of these unique signals from the PrediXcan TWAS appeared only in one or two tissues (92.5% for HDL, 98.2% for LDL and 100% for T2D).

With all these various considerations, SWAM appears to improve TWAS power for a given tissue, although ultimately may yield fewer signals compared to comprehensive tissue scans using PrediXcan or MultiXcan. While SWAM outperforms other methods in terms of imputation accuracy, there may not be a clear-cut winner in terms of performance in TWAS. The best approach to use will likely depend on the needs of the researcher, and each approach may provide different yet complementary insights into understanding the biological mechanisms from these association studies.

## Discussion

The transcriptome serves as an intermediate phenotype linking genetic variants to complex traits. Association studies between traits and gene expression, when used in conjunction with GWAS, provide additional insight into the biological mechanisms of complex traits. Imputation of gene expression in the context of transcriptome wide association studies is a promising approach to understanding the connection between our genes and many traits. Yet, there are still many challenges that arise when performing association studies with imputed expression. Current tissue-specific imputation models are trained using data obtained from their respective tissues, which can vary greatly in data quality and sample size. As such, there is a great deal of variability among tissues in the imputation accuracy of tissue-specific gene expression levels. For example, PrediXcan was able to significantly impute only 2086 vagina-specific genes, while it discovered 8171 genes specific to the tibial nerve tissue. Furthermore, the imputation accuracy of significant genes within a tissue are also highly variable, with some genes such as ERAP2 having very high (>80% of variation explained by eQTLs) imputability and other genes (~1% of variation explained by eQTLs) with low imputability.

In this paper we developed SWAM, a method that determines the level of eQTL sharing between tissues and uses the shared information from other tissues to improve the imputation accuracy for the target tissue. By simultaneously examining the relatedness of multiple tissues, SWAM in essence increases the effective sample size of imputation models. Using GEUVADIS LCL data, we compared SWAM to single-tissue approaches. We found that our multi-tissue approach, in addition to increasing the number of significantly imputable genes for each tissue, also improved the overall imputation accuracy for genes that were already significantly imputable using PrediXcan. We improved the power of TWAS by running a SWAM-adapted version of MetaXcan for various traits, finding an increased number of significant transcriptome-trait associations, even when correcting for the larger number of genes imputed.

Although SWAM provides a substantial improvement for the number of significantly imputable genes for many tissues and generally increases power for TWAS, there are some shortcomings and caveats to consider with the approach. It is important to note that unlike PrediXcan, SWAM does not actually perform model training or eQTL discovery. Instead, it evaluates the efficacy of various single-tissue imputation models (in this case, the GTEx tissues) and assigns weights to the models based on their relatedness to the target tissue. Therefore, for SWAM to work, there must already be a database of imputation models that it can use to derive the multi-tissue weighting. Because we are utilizing existing imputation models, we acknowledge that there will be cases where the SWAM imputation accuracy could be similar or worse to the single-tissue imputation, especially if the gene has shared eQTLs across many tissues or if the single-tissue imputation model was already performing well. The improvement observed in our validations and TWAS are an overall trend, and as with any analysis, interpretation of any specific results should be approached with caution. Furthermore, the improvement for any given gene has an upper limit which is dependent on the pool of single tissue models available. There may be tissues that have very few relevant other tissues to draw information from. For any given gene within the target tissue, SWAM automatically assigns weights of non-relevant tissues to zero based on a threshold. However, for the purposes of our study, the threshold was tuned to be more lenient, allowing for more tissues to be included in the imputation of each gene’s expression levels. A more lenient threshold will yield more genes, but a lower sensitivity to the target tissue. A stricter threshold will provide imputed expressions that are more specific to the target tissue but will provide imputation for fewer genes and may reduce imputation accuracy in some genes. Optimal tuning of this threshold may depend on the target tissue, and the goals of the analysis. Further work could help determine the ideal way to tune these thresholds, perhaps using a different threshold depending on the gene and tissue in question.

Next, our empirical validation of imputation accuracy was tested on European individuals (344 samples from GEUVADIS) and thus SWAM’s performance with other populations has not yet been determined. A future direction of research could be to examine whether a single model derived from mixed populations would represent each of the populations accurately, or if a different model should be trained on each population separately. Currently, evidence suggests that training from the correct ancestry group is the ideal approach for population-specific imputation [32], which emphasizes the importance of reference panel resources derived from a wide array of ancestries. Alternative approaches could be to leverage trans-ancestry correlation, which has been shown to increase predictive R^2^ in the context of polygenic risk scores [33].

Finally, while SWAM improved the number of association signals for any given tissue in TWAS compared to UTMOST and single tissue PrediXcan, aggregation of signals (MultiXcan/combining PrediXcan signals) suggest that other approaches may yield more unique signals. It is unclear which approach is preferable in this scenario, and the answer may depend on unraveling the causality of association signals. Recently, there have been a number of publications which have addressed this issue, such as PTWAS which uses instrumental variables (IVs) to investigate the causal relationship between expression levels and complex traits [26], or phenomeXcan, which integrates GWAS and gene expression and regulation data to identify likely causal pathways [34]. Future directions could include using IVs or functional annotation to interpret TWAS signals.

To conclude, we propose a novel method for gene expression imputation, which extends already established single-tissue imputation models into a multi-tissue setting. By combining information from multiple models, we were able to increase overall tissue-specific imputation accuracy for many genes and increase power for transcriptome-wide association studies.

## Materials and Methods

### SWAM Notation and Framework

Our framework for *SWAM* is designed to find the optimal linear combination of imputed expression levels from multiple tissues and datasets. For simplicity, we will denote each (tissue, dataset) combination as a source. We assume there are *K* imputation models from individual sources, with each model indexed as *j* ∈ (1, . ., *K*). We also denote *r* ∈ {1, …, *K*} to represent the index of the reference source. The inputs for *SWAM* are: (1) *f*_j_(·) – the single-source imputation models and (2) **Y**_*r*_ and **X**_*r*_ – the individual-level gene expression measurements and genotypes for the reference source. For each gene *g*, let 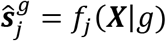 be imputed expression from a single source. Then we can represent any linearly combined multi-tissue imputed expression 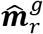 as

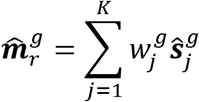

where 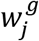 is the weight contributed by *j*-th source. *SWAM* learns 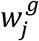 by leveraging individual-429 level data from the reference source as we describe later.

### Multi-tissue methods using naïve average or best-tissue

There are two heuristic approaches to impute expressions from multiple sources - *naïve average* and *best tissue*. *Naïve average* defines weights uniformly as 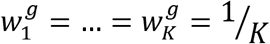 For *best tissue,* the weights are defined as a dichotomous variable:

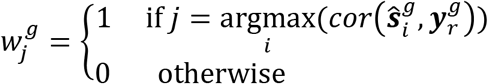

where 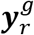 represents the individual-level expression measurements of the reference source.

### Smartly Weighted Average across Multiple Datasets (SWAM)

Here we describe how SWAM calculates optimal 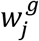, whose derivation is shown in the Supplementary Text. It is important to note that *SWAM* works ideally when the tissue type intended to be imputed matches to the tissue types of the reference source. We define 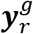 as the *n* × 1 vector of individual-level expression measurements for the reference source, and as before, **X**_*r*_ to be the corresponding *n* × *m* matrix of individual-level genotypes. The first step is to impute expression using each of the *K* models using the reference genotypes. Thus, we obtain *K* sets of imputed expressions, 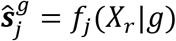, with each being a single-source imputation for the samples in the reference data. The weights for *SWAM* are given by

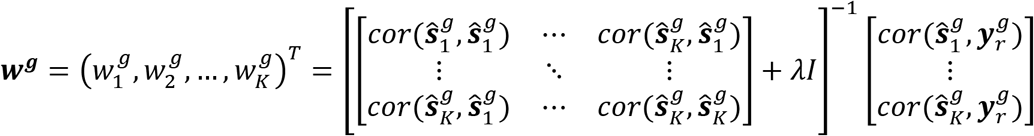

Here, the correlation matrix account for the similarity between the imputation models, and the vector containing the entries 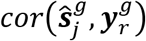account for the empirical similarity of imputed expressions from each model to the measured expressions in the reference source. When *j* = *t*, because 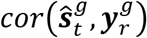 will be prone to overfitting, we replace this value to a 5-fold cross-validated correlation instead, which is available from PrediXcan output. Finally, λ*I* acts to regularize the weights, providing numerical stability for the inversion of the covariance matrix. The calibration of λ is further discussed in the Supplementary Text.

### Simulations

Our simulation study sought to examine SWAM’s ability to detect the correct shared components between related tissues across a wide spectrum of parameter settings. We compared *SWAM* with *naïve average, best tissue* and *single tissue* approaches. For each simulation, we independently generate individual-level genotypes and expression multiple tissues. For the reference set, we simulated *X*_*r*_, an *n*_r_ × *m* genotype where *n*_*r*_ is the number of individuals and *m* the number of SNPs. In our simple simulation, we assume that each SNP is independent, with non-reference allele frequency (AF) distributed with *Beta(1,3)*. The genotypes were simulated using a binomial distribution based off the AF. To simulate multi-tissue expressions, for each tissue *j* ∈ (1, . ., *K*) we specific effect sizes **β**_**j**_, to simulate expressions **y**_**j**_ = *X*_r_**β**_**j**_ + ɛ_j_. For reference tissue (i.e. *j* = *r*), we assume two causal SNPs with nonzero elements in **β**_**j**_, where one SNP is expected to explain tissue-specific heritability 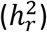 for the reference tissue and the other SNP explains the cross-tissue heritability 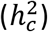, summing up to total heritability 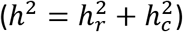. Other tissues (i.e. *j* ≠ *r*) were divided into “related tissues” and “independent tissues”. For related issues, **β**_**j**_ had only one non-zero values corresponding to cross-tissue heritability 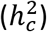. For independent tissues, all **β**_**j**_ had zero values. Finally, we generated another set of validation genotypes matrix *X*_*v*_ with size *n*_*v*_ × *m*, and the validation expressions (**y**_*v*_ = X_*v*_**β**_***r***_ + ɛ_*v*_) of reference tissue using the same settings to use for evaluation.

We then trained tissue-specific imputation models *f*_j_(.), *j* ∈ (1, …, *K*) by applying an elastic-net model (using *glmnet* R package [35]) for each pair of *X*_r_ and **y**_***j***_. The tuning parameters for elastic net were determined via a five-fold cross-validation technique. Using **y**_***r***_, *X*_r_ and *f*_i_(.), we obtained *naïve average, best tissue* and SWAM models as detailed in the framework and weights section. To calculate the proportion of imputable genes, we performed linear regression between **y**_v_ and the imputed expression from genotypes *X*_v_ using the different methods to obtain a p-value.

Each simulation was repeated for 1,000 times in each setting. We varied parameters to evaluate their impact on the performance of each method. We varied *h*^2^ ∈ {0, 0.1, …,1} (default 0.1), 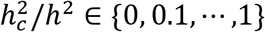 (default 0.5), *K* ∈ {2, 4, 6, 8, 10, 20, 30, 40, 50} (default 10), fraction of independent tissues ranging {0, 0.1, …,0.8} (default 0.5), *n*_r_ ∈ {50,100, …,500} (default 200), and the p-value threshold ranging {10^−6^, 10^−5^, …, 0.01, 0.05, 0.1} (default 0.05). Throughout all simulations, *m* = 35, *n*_v_ = 200 were used.

### Input Datasets: Genotypes, Expressions, and Imputation Models

In our experiments with real datasets, we leveraged multiple published datasets where genotypes, expressions, and imputation models are available to evaluate the performance of SWAM and other methods in various settings. Specifically, we used the GEUVADIS LCL [29] genotypes and expressions as a validation dataset. We used GTEx data [14] [24] and PredictDB [2] to build multi-tissue imputation models. To demonstrate the ability to SWAM to incorporate multiple datasets, we used DGN [11] dataset as well as multiple versions of GTEx datasets.

### Multi-tissue transcriptomic profiles and imputation models from the GTEx project

To build multi-tissue imputation models using *SWAM, UTMOST, naïve average, and best tissue methods*, we used single-tissue imputation models, individual-level genotypes, and expressions obtained from the GTEx consortium. Single-tissue imputation models were downloaded from the PredictDB (http://predictdb.org/) repository for GTEx versions 6, 7 and 8 (44, 48 and 49 tissues respectively) [3] [14] [24], which were trained using PrediXcan’s elastic net methods. Individual-level genotypes and expression levels were only used for the reference tissue (e.g. EBV-transformed lymphocytes) which is deemed to be the closest to the validation data (e.g. GEUVADIS LCL), using GTEx version 6.

When evaluating multi-tissue imputation models within a single dataset, we used GTEx version 6. When evaluating imputation models across multiple tissues and multiple datasets, we used various combinations of GTEx versions to evaluate the benefit of multiple imputation models trained from overlapping datasets. When training across different datasets, genes were matched by ensemble ID, ignoring version numbers. In addition to training SWAM, we also used the *single tissue* PredictDB imputation models as a basis for comparison with our method.

### Validation dataset from the GEUVADIS study

We used individual-level genotypes and expression levels from lymphoblastoid cell lines (LCL) from the GEUVADIS consortium only to evaluate various methods after imputing expression levels with models built from other datasets. Each imputation model was evaluated by applying the model to GEUVADIS genotypes to impute individual expression levels, and by calculating the correlation between the imputed and measured expressions. We focused on 344 European individuals where genotypes and normalized expressions (from RNA-seq) are available, with comparable linkage disequilibrium (LD) structure to GTEx and DGN datasets.

### Imputation models from Depression Genes Network

We also downloaded the imputation model trained using the 922 whole blood transcriptomes from the Depression Genes Network (DGN) via PredictDB. DGN was evaluated as a single-tissue imputation model. It was also used in the evaluation of multi-dataset imputation models when DGN is combined with various versions of GTEx imputation models.

### Imputation models from UTMOST

We compared our methods to *UTMOST*, another multi-tissue approach for expression imputation [24]. The *UTMOST* imputation models were jointly trained across 44 tissues from GTEx version 6 and were downloaded from their published online repository (https://github.com/Joker-Jerome/UTMOST). We applied the imputation model targeted for EBV-transformed lymphocytes when evaluating the imputation accuracy with the GEUVADIS LCL expression.

### Evaluating imputation accuracy with GEUVADIS measured expression

We evaluated the accuracy of various imputation models by comparing imputed expressions from individual-level genotypes with the measured expression from GEUVADIS LCLs. Individual-level expression were imputed across 344 European GEUVADIS samples using various single-tissue, multi-tissue/multi-dataset methods to calculate the correlation with the normalized measured expression from GEUVADIS LCL. The correlation between imputed and measured expressions were calculated using spearman correlation and a one-sided p-value was evaluated by converting the correlation coefficients into t-statistics. Genes were considered “significantly imputable” if the Benjamini-Hochberg false discovery rate (FDR) was less than 0.05. This procedure was applied across all genes within each method, with the counts being tabulated.

### Comparing single-tissue and multi-tissue imputation models within a single dataset

With these results, we first focused on comparing the imputation accuracy of SWAM with other methods using GTEx v6. We compared *SWAM-LCL* (SWAM using GTEx EBV-transformed lymphocytes as reference), every *single tissue* imputation model from PredictDB, *UTMOST-LCL* (*UTMOST* using GTEx EBV-transformed lymphocytes as reference), *naïve average*, and *best tissue* methods. We focused on evaluation using GTEx v6 models where *UTMOST* models were available. We also focused on genes included in the Consensus Coding Sequence Project (CCDS) [36] to minimize the discrepancy between imputation models.

To keep a fair comparison with *UTMOST* and the *single tissue* methods, we restricted the set of genes to those that have at least one eQTL in any *single tissue* models from PredictDB and also in any *UTMOST* models across all reference tissues.

### Evaluating multi-tissue imputation models across multiple datasets

Our second comparison was conducted to examine the effect of integrating multiple imputation models trained from heterogeneous datasets into SWAM. Here, we used various combinations of GTEx and DGN resources to derive multi-tissue/multi-dataset models, such as combining GTEx v6 with DGN data, or combining GTEx v6, v7 and v8 altogether. For this analysis, the gene list was restricted to genes that were included in all three of the v6, v7 and v8 datasets in terms of Ensemble IDs.

### Evaluation of SWAM in transcriptome-wide association studies (TWAS)

To evaluate our method in the context of TWAS, we used MetaXcan [30], which infers TWAS results from GWAS summary statistics. We focused on the HDL and LDL traits from Global Lipids Genetics Consortium (GLGC) [37] and Type-2 Diabetes (T2D) from the DIAGRAM consortium [38]. For this analysis, we generated SWAM imputation models targeting each of the 44 tissues from GTEx version 6. We used MetaXcan to infer the TWAS results for each of these tissues and applied a Bonferroni correction with false-positive rate of 0.05 based on the number of genes tested. We repeated this with all 44 UTMOST models as well as all 44 PrediXcan *single tissue* models.

We also compared our method with S-MultiXcan [25], a recently published extension of MetaXcan which uses a principal components regression to conduct trait-expression association with multiple tissues.

## Software Availability

The software package for SWAM is available at https://github.com/aeyliu/swam, including the datasets used in this manuscript.

## List of Figures

**Figure 1 – overview of SWAM method.**

This figure demonstrates the training of the imputation model using the reference data. The inputs required for SWAM are a set of reference genotypes with sample matched measured expression, and the multiple imputation models to be included. The list of multiple imputation models must also include a model derived from the reference data, which can be done via prediXcan. SWAM uses these models to impute tissue-specific expression levels from the reference genotypes. These imputed expression sets are then compared with the measured expression of the reference set. The weights are calculated based on the similarity between the measured and imputed expression and the covariance structure of tissues. For full details, see the methods section.

**Figure 2 – simulation study comparing SWAM with naïve average, best tissue and single tissue methods.**

We ran each simulation 10,000 times, with the following default settings: 10 total tissues (1 target, 4 relevant, 5 irrelevant), 100 SNPs (2 per tissue), 10% genetic heritability, 50% shared heritability between relevant tissues. In addition, the sample size of the target tissue was 100 individuals, and the remaining tissues had 200 individuals. This was done to emphasize the importance of integrating information from other tissues when the quality of the target tissue model is limited. In panel (A), we varied the number of relevant tissues, from 0 to 10. Panel (B) shows the improvement when the total number of tissues is increased, with the number of irrelevant tissues fixed at 50% of the total. Panel (C) shows the effects of changing the shared heritability for the relevant tissues. We note here, that each tissue has 2 causal SNPS – for the relevant tissues, 1 of these causal SNPS is shared with the target tissue while the other is independent of all simulated tissues. Panel (D) shows the performance of the approaches for different levels of genetic heritability. This simulation demonstrates the range of heritability that we would expect to see the most improvement. Empirically, we do notice the same trend seen here, as SWAM performs similarly the single tissue model when the cross-validated R-squared is high. Panel (E) shows the effects of target tissue sample size. The x-axis pertains to the sample size of the target tissue only, and all other tissues were fixed at 200 individuals. Finally, panel (F) shows the performance of the methods at different p-value thresholds, using the default simulation settings.

**Figure 3 – Empirical validation of SWAM using lymphoblastoid-cell line data from GEUVADIS consortium.**

We used our LCL-targeted SWAM model to impute expression levels based on the genotypes of 344 European samples. We then calculated the concordance between imputed expression and measured LCL expression. We repeated this for all of the other methods mentioned here. (A) shows the performance of SWAM against the single-tissue models from 44 tissue-specific predictDB models derived from GTEx version 6. In (B), we derived various SWAM models using every combination of the following: 1) all GTEx v6 tissues, 2) all GTEx v7 tissues, 3) all GTEx v8 tissues, and 4) Depression Gene Network (DGN) single tissue whole blood model from predictDB. Here, we also included the UTMOST LCL model, naïve average and best tissue models, all derived from GTEx v6.

**Figure 4 – TWAS on LDL trait targeting liver using SWAM, UTMOST and PrediXcan models.**

TWAS was performed using metaXcan on the LDL trait from the Global Lipids Genetics Consortium (GLGC) GWA analysis. For a consistent comparison, the SWAM and UTMOST models were derived from GTEx version 6 tissues, and the prediXcan model used was GTEx v6 liver. The number of associations were: 74, 69 and 19 for SWAM, UTMOST and prediXcan respectively. P-values were truncated at 10^−20^ in these plots.

## Supplementary Materials

### Derivation of weights for SWAM

In this section we derive the equation for the weights in SWAM. We wish to impute expression for a reference sample of *N* individuals with genotypes *X*_*r*_ and measured tissue-specific expression **y**_***r***_. Suppose we have single tissue imputation models for *K* tissues, with *r* ∈ {1, …, *K*}. For each gene *g*, we obtain a set of *K* imputed expression levels 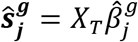, with *j* ∈ (1, …, *K*). Dropping the superscript *g* for convenience, we define *w* = (*w*_l_, *w*_2_, …, *w*_K_)′ be the set of weights corresponding to each of the tissues. The SWAM estimator is thus:

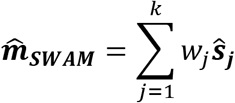

For further convenience, we denote 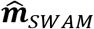 as 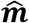. Then, for each gene separately, the values for *W* are determined by minimizing the expression:

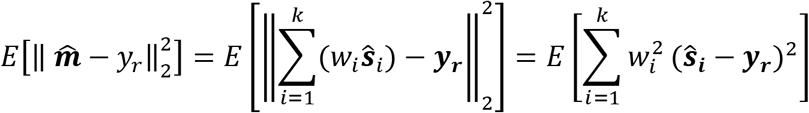

Without loss of generality, we set the constraint 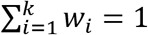. The objective function to be minimized is

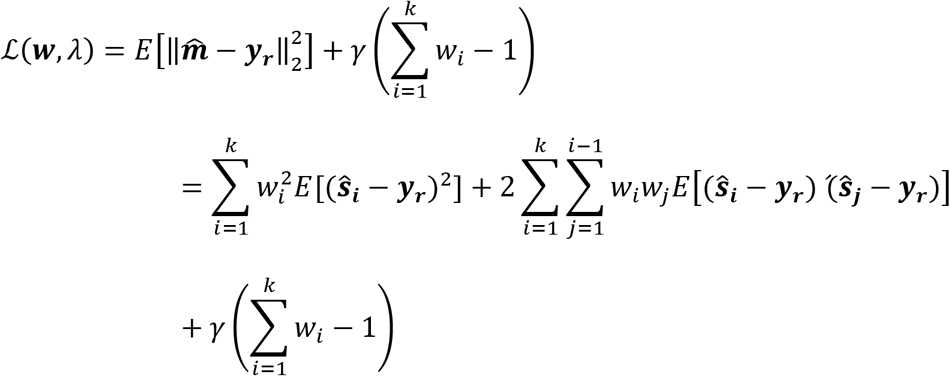

The gradient of 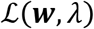 is

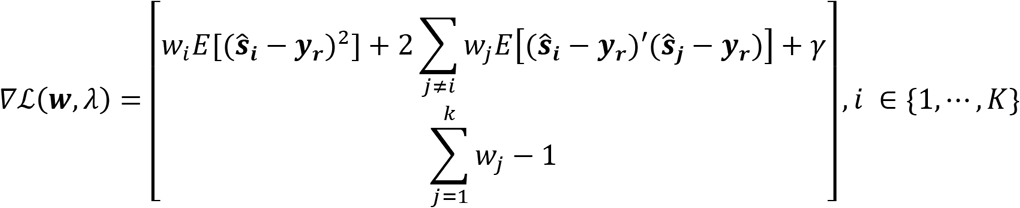

Solving this system of equations, we obtain the optimal weighting minimizing the expected MSE across single tissue imputed expressions as

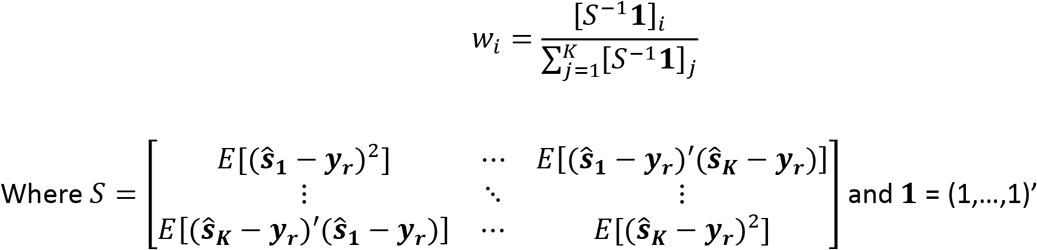

### Regularization of weights

The weights derived in the previous section provide an optimal solution to the expression 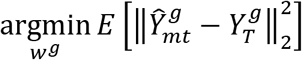. In the scenario in which the tissues are highly correlated with each other, the matrix 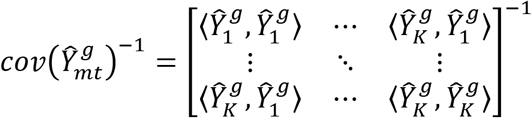 is numerically unstable as the columns of 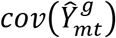) are no longer linearly independent. This can lead to high weights assigned to irrelevant tissues and lower weights for relevant tissues. Furthermore, this may result in weights that are over-fitted to the noise of the data. To correct for this, we added a diagonal matrix, λ*I* prior to inverting the matrix 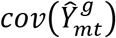, giving us the solution 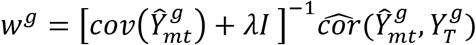. To choose the correct value of λ, we tested the imputation accuracy of 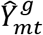 in our validation test set for a large range of λ. We found that imputation accuracy was low when λ = 0, likely due to overfitted and the amplification of noise. Larger values of λ yielded better results but ignored the correlation structure between tissues. We found empirically that λ = 3 provided the best results (this value depends highly on the scale and normalization of the data).

### Application of SWAM to other target tissues

Throughout our work we primarily used the LCL tissue from GTEx version 6 as our target tissue for application of SWAM. In addition to producing SWAM-LCL models, we also generated models targeting each of the 44 GTEx v6 tissues. Supplementary Figure 3 displays the heatmap of weight contribution towards each of the tissues. The rows correspond to the SWAM model for each tissue type, and the color intensity of the columns show the contribution of each tissue towards the targeted tissue (number of times the tissue contributed the highest weight). Overall, we observe clustering that appears to separate the tissue types quite well. For example, brain tissues are primarily getting high weights from other brain tissues while receiving low weights from all other tissue types. This heatmap provides evidence of SWAM being able to capture tissue-specific signals.

---

**Supplementary Figure 1 – Using SWAM to impute expression and conduct TWAS**

The first panel shows how SWAM can be used to impute expression levels via prediXcan, while the second panel shows the required inputs to conduct TWAS via metaXcan.

**Supplementary Figure 2.**
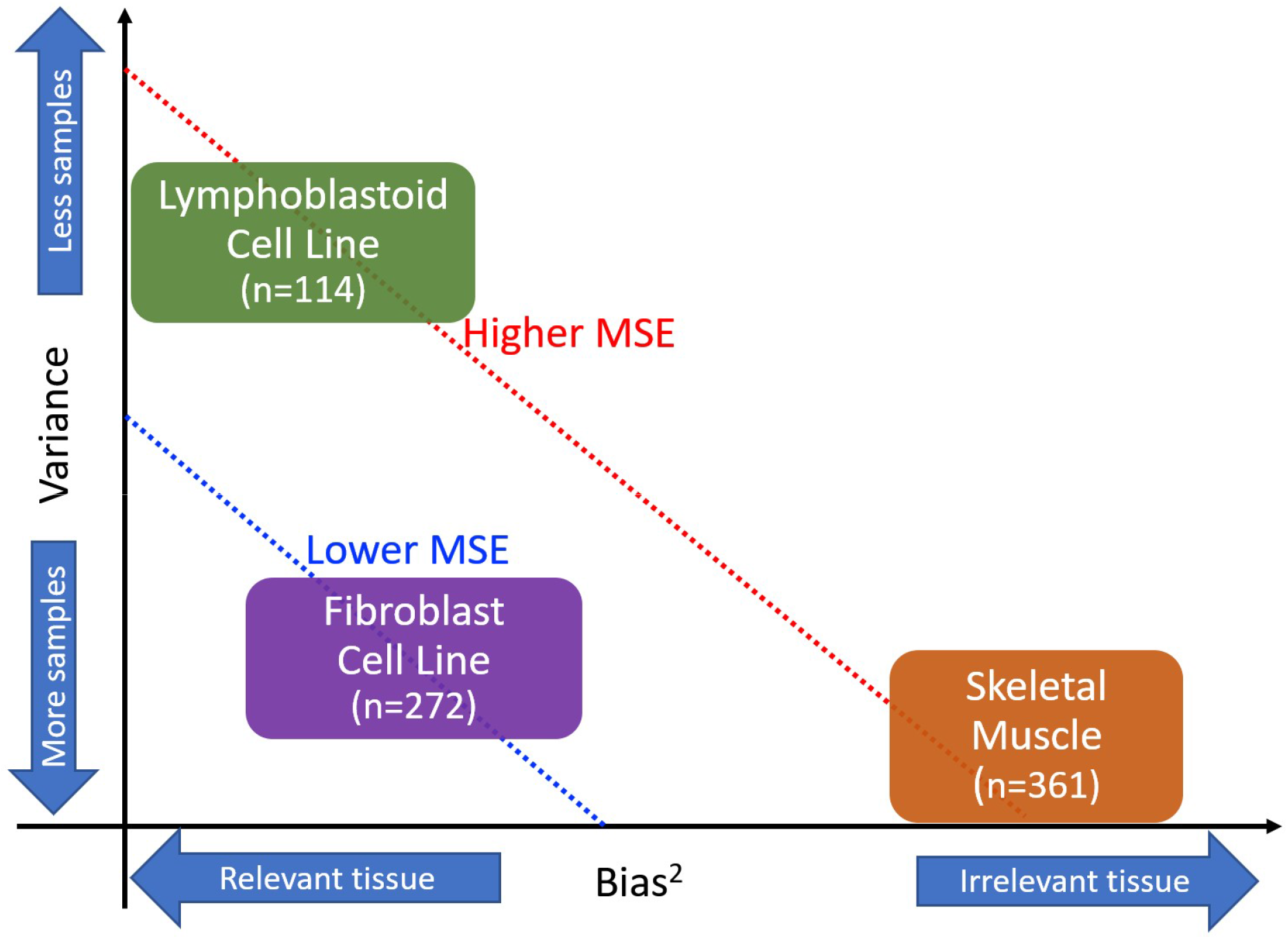
Bias-variance tradeoff for other tissues. The principal behind SWAM is it considers the bias-variance tradeoff for each tissue, and assigns higher weights to tissues that reduce MSE. In this example, tissues such as Skeletal Muscle have a high sample size (and therefore lower variance) but may be biased as they are not the relevant tissue to the tissue of interest (in this case LCL). Other tissues such as Fibroblasts may have a lower sample size but compensate by having low bias (high relevance to tissue of interest) and will contribute more weight.

**Supplementary Figure 3–.**
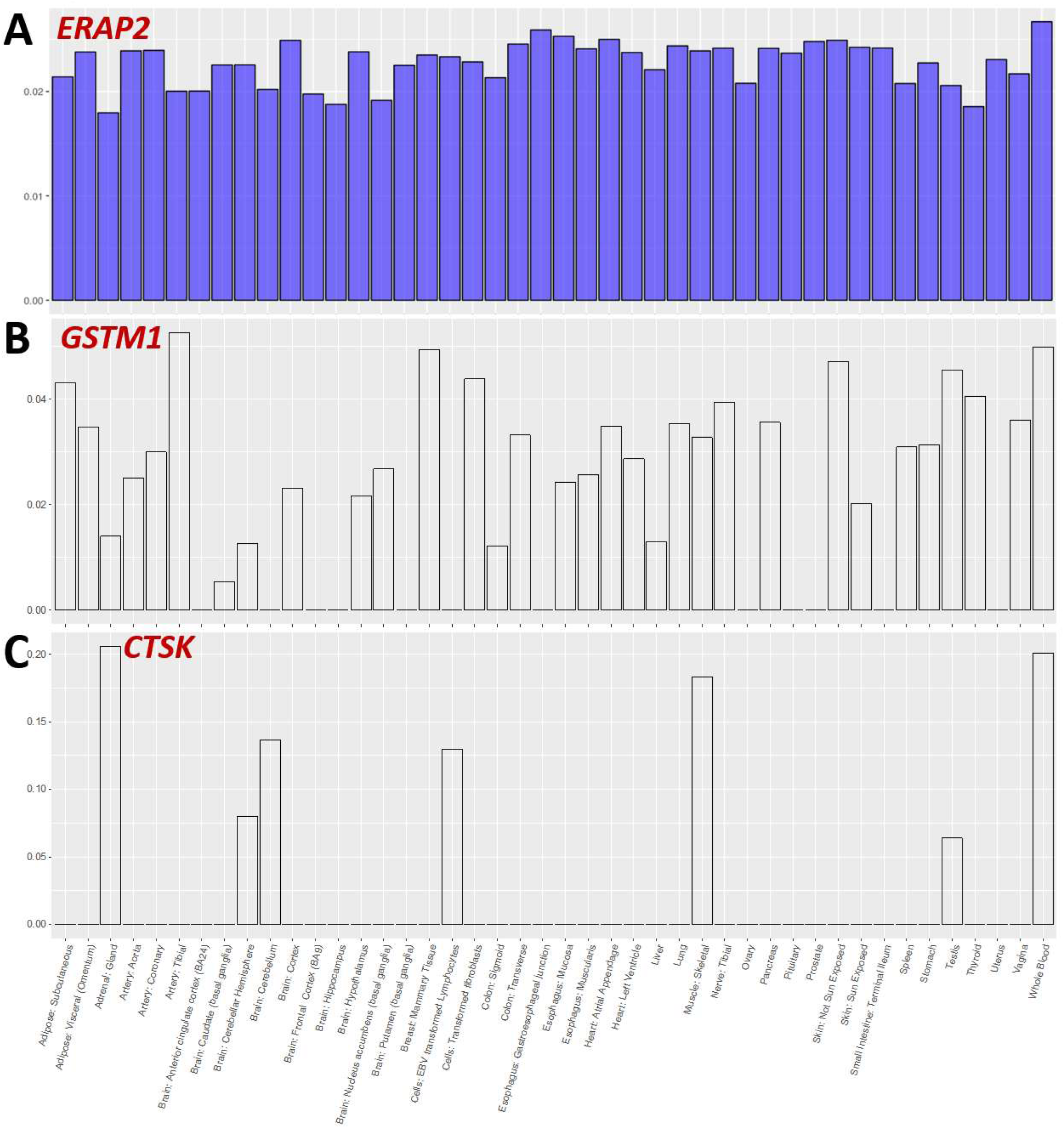
The distribution of weights for SWAM for three selected genes. (A) shows the ERAP 2 gene, which had a single tissue r^2^ = 0.801, while the SWAM model had r^2^ = 0.795. (B) depicts are scenario where SWAM is able to leverage information from other tissues to make up for the relatively lower quality of the target tissue – here the single tissue model gave r^2^ = 0.368 while SWAM increased the accuracy to r^2^ = 0.741. (C) shows an example where the eQTLs are highly tissue specific. Here, SWAM improved the single tissue accuracy from r^2^ = 0.111 to r^2^ = 0.447.

**Supplementary Figure 4.**
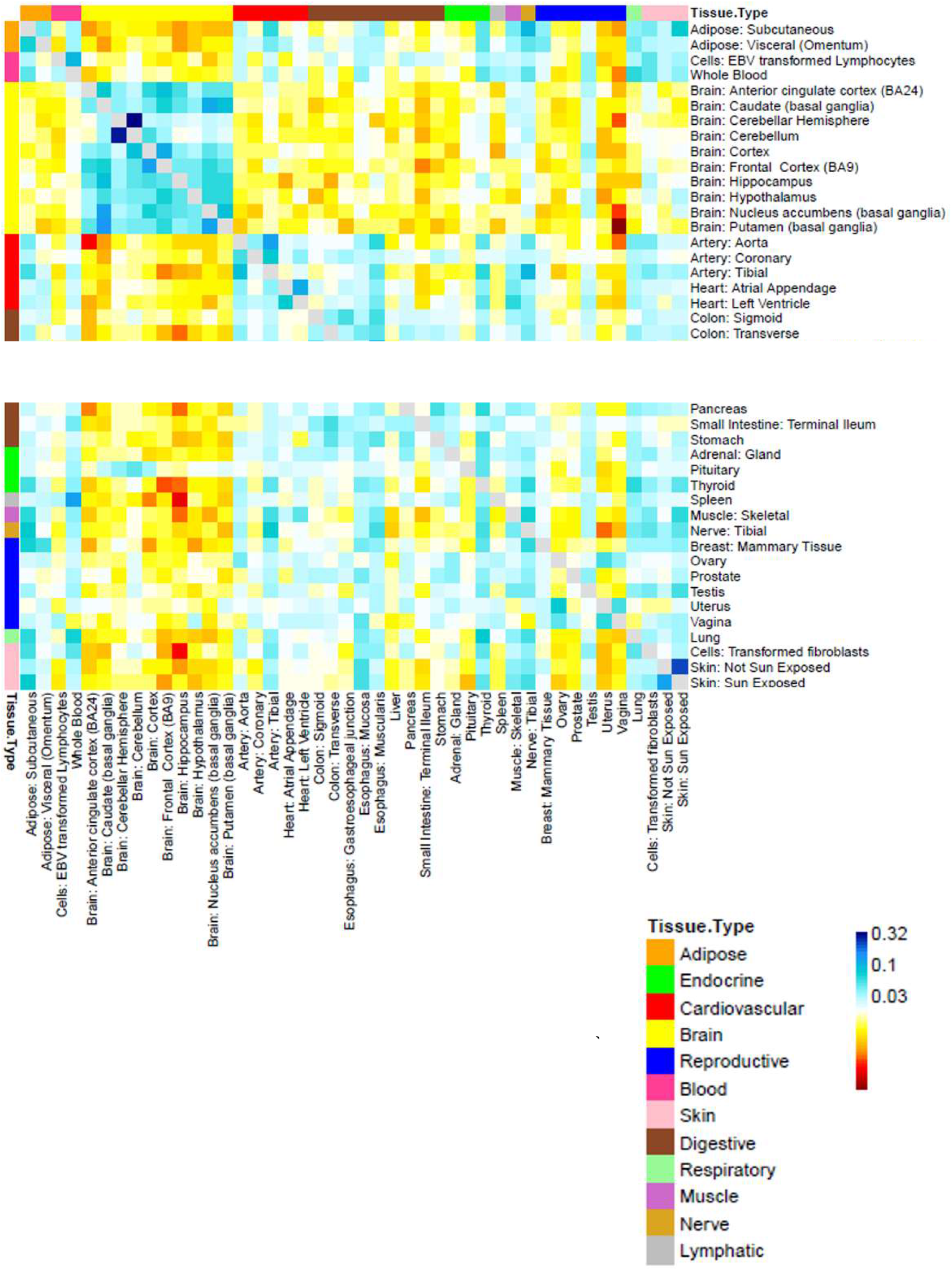
distribution of SWAM weights in imputation models for all 44 GTEx v6 tissues. Here, we used SWAM to derive multi-tissue imputation models for all 44 GTEx v6 tissues. Each cell in this heatmap depict the number of times each tissue contributed the highest weight to the target tissue. Here, the rows correspond to the target tissue and the columns correspond to the weight contribution of each tissue. For the sake of clarity, the diagonal values were not included as they were consistently much higher than the remaining elements of the matrix.

**Supplementary Figure 5.**
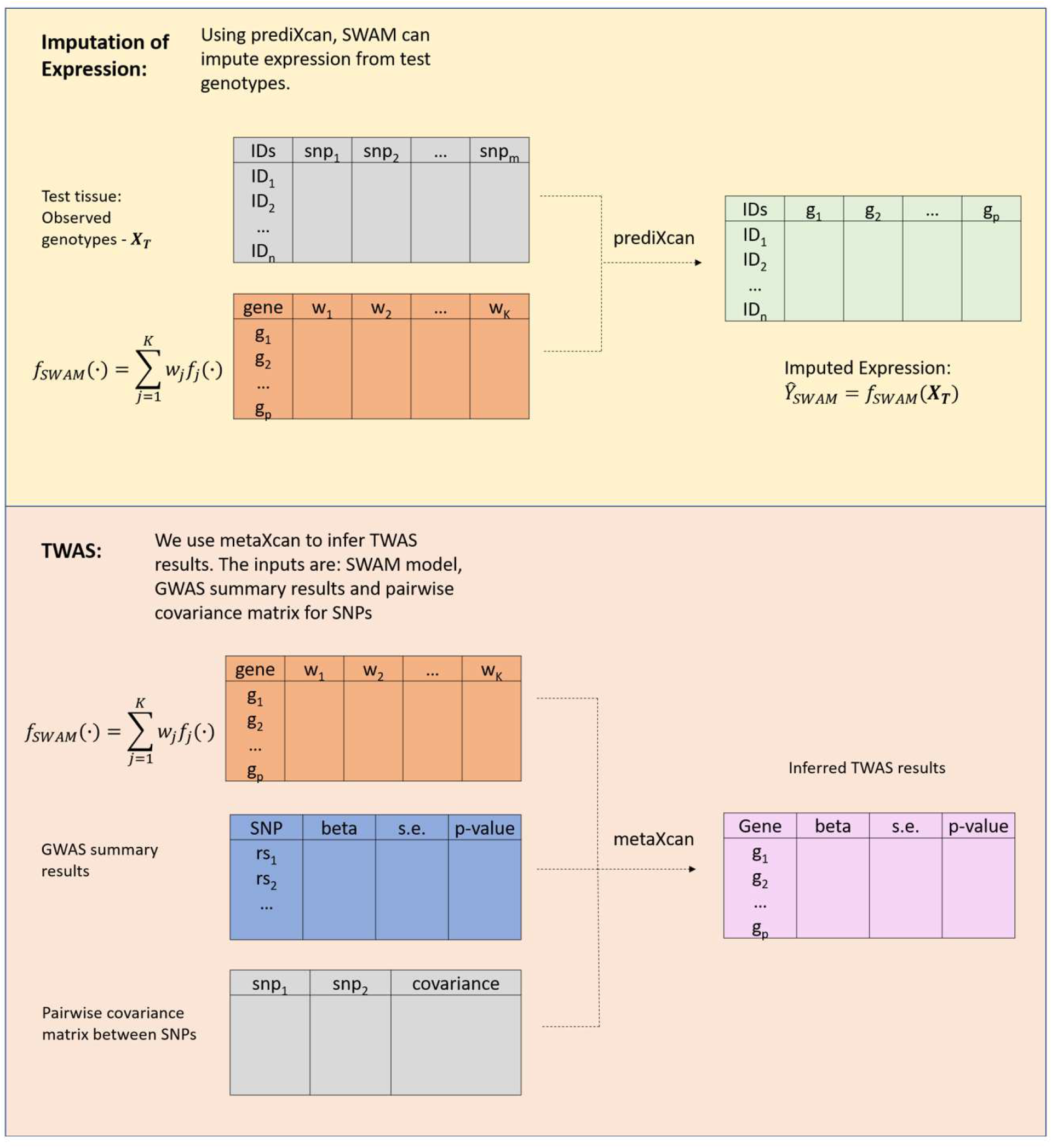
Using SWAM to impute expression and conduct TWAS. The first panel shows how SWAM can be used to impute expression levels via prediXcan, while the second panel shows the required inputs to conduct TWAS via metaXcan.

**Supplementary Table 1 – GTEx version 6 comparisons of single-tissue and multi-tissue imputation models using GEUVADIS LCL RNA-Seq expression as validation.**

*Counts (B-H counts) are based on Benjamini-Hochberg procedure false discovery rate of 0.05. The last column displays the number of counts at p-value threshold 0.05 (without any corrections)*

**Supplementary Table 2– Comparison of all multi-tissue methods**

*We applied SWAM to all combinations of GTEx and DGN resources. For the GTEx resources, we always used every tissue available. In version 6, this comprised of 44 tissues. For version 7, there were 48 tissues and version 8 contained 49 tissues. For the sake of consistency, our target tissue for each of these combinations was GTEx v6 LCL.*

**Supplementary Table 3 – comparison of GTEx v7/v8 single tissue models versus GEUVADIS LCL**

*We also compared every prediXcan model derived from GTEx version 7 and version 8 tissues, and tested imputation accuracy against GEUVADIS LCL measured expression levels. Surprisingly, despite the increase in sample size, the LCL tissue from v8 performed worse than its version 7 counterpart. The number of tissues outperforming LCL in both v7 and v8 highlight the opportunity to leverage information from other tissues to improve imputation accuracy for under-powered tissues.*

**Supplementary Table 4– TWAS association signals for SWAM**

*We used SWAM to derive an tissue-specific model for every GTEx version 6 tissue, and used these models as inputs to metaXcan to infer TWAS results. As mentioned in the methods section, the HDL and LDL traits were from Global Lipids Genetics Consortium (GLGC) and Type-2 Diabetes (T2D) from the DIAGRAM consortium.*

**Supplementary Table 5 – TWAS association signals for UTMOST**

*These models were also derived from GTEx version 6 tissues using the UTMOST method. Models were downloaded from https://github.com/Joker-Jerome/UTMOST*

**Supplementary Table 6 – TWAS association signals for prediXcan (single-tissue)**

*TWAS results via metaXcan using prediXcan single tissue models derived from GTEx version 6 tissues*

**Supplementary Table 7 –.** GTEx version 6 comparisons of single-tissue and multi-tissue imputation models using GEUVADIS LCL RNA-Seq expression as validation. *Counts (B-H counts) are based on Benjamini-Hochberg procedure false discovery rate of 0.05. The last column displays the number of counts at p-value threshold 0.05 (without any corrections)*

| Method | Total # genes | Genes with FDR < 0.05 | P-value threshold for FDR=0.05 | Genes with p-value < 0.05 |
| --- | --- | --- | --- | --- |
| DGN Whole Blood | 13213 | 2390 | 0.009018 | 3450 |
| Adipose Subcutaneous | 13213 | 1500 | 0.005667 | 2203 |
| Adipose Visceral Omentum | 13213 | 963 | 0.003619 | 1432 |
| Adrenal Gland | 13213 | 735 | 0.002755 | 1151 |
| Artery Aorta | 13213 | 1177 | 0.004426 | 1783 |
| Artery Coronary | 13213 | 616 | 0.00231 | 926 |
| Artery Tibial | 13213 | 1439 | 0.005399 | 2124 |
| Brain Anterior cingulate cortex BA24 | 13213 | 322 | 0.00117 | 580 |
| Brain Caudate basal ganglia | 13213 | 528 | 0.001979 | 895 |
| Brain Cerebellar Hemisphere | 13213 | 575 | 0.002172 | 1002 |
| Brain Cerebellum | 13213 | 661 | 0.002498 | 1144 |
| Brain Cortex | 13213 | 511 | 0.00193 | 878 |
| Brain Frontal Cortex BA9 | 13213 | 456 | 0.001699 | 770 |
| Brain Hippocampus | 13213 | 336 | 0.001254 | 572 |
| Brain Hypothalamus | 13213 | 339 | 0.001264 | 554 |
| Brain Nucleus accumbens basal ganglia | 13213 | 454 | 0.001709 | 755 |
| Brain Putamen basal ganglia | 13213 | 391 | 0.001461 | 655 |
| Breast Mammary Tissue | 13213 | 933 | 0.003455 | 1430 |
| Cells EBV-transformed lymphocytes | 13213 | 1552 | 0.005845 | 1943 |
| Cells Transformed fibroblasts | 13213 | 1690 | 0.00635 | 2451 |
| Colon Sigmoid | 13213 | 680 | 0.002391 | 1052 |
| Colon Transverse | 13213 | 1013 | 0.003803 | 1536 |
| Esophagus Gastroesophageal Junction | 13213 | 716 | 0.002689 | 1101 |
| Esophagus Mucosa | 13213 | 1469 | 0.00553 | 2148 |
| Esophagus Muscularis | 13213 | 1272 | 0.004808 | 1959 |
| Heart Atrial Appendage | 13213 | 849 | 0.003167 | 1323 |
| Heart Left Ventricle | 13213 | 942 | 0.003537 | 1451 |
| Liver | 13213 | 427 | 0.001547 | 719 |
| Lung | 13213 | 1355 | 0.005122 | 1985 |
| Muscle Skeletal | 13213 | 1197 | 0.004454 | 1876 |
| Nerve Tibial | 13213 | 1450 | 0.005483 | 2186 |
| Ovary | 13213 | 417 | 0.001566 | 689 |
| Pancreas | 13213 | 911 | 0.003426 | 1406 |
| Pituitary | 13213 | 496 | 0.001875 | 808 |
| Prostate | 13213 | 398 | 0.001421 | 629 |
| Skin Not Sun Exposed Suprapubic | 13213 | 1063 | 0.003997 | 1619 |
| Skin Sun Exposed Lower leg | 13213 | 1463 | 0.005489 | 2145 |
| Small Intestine Terminal Ileum | 13213 | 450 | 0.001694 | 735 |
| Spleen | 13213 | 759 | 0.002821 | 1166 |
| Stomach | 13213 | 878 | 0.003291 | 1350 |
| Testis | 13213 | 956 | 0.003607 | 1563 |
| Thyroid | 13213 | 1454 | 0.005469 | 2208 |
| Uterus | 13213 | 297 | 0.001071 | 521 |
| Vagina | 13213 | 307 | 0.001149 | 511 |
| Whole Blood | 13213 | 1427 | 0.005343 | 2106 |
| SWAM-LCL v6 | 13213 | 3040 | 0.01145 | 4148 |
| NAIVE AVERAGE V6 | 13213 | 2666 | 0.010066 | 3830 |
| BEST TISSUE V6 | 13213 | 2493 | 0.009394 | 3663 |
| UTMOST-LCL v6 | 13213 | 2238 | 0.008466 | 3185 |

**Supplementary Table 8 –.** Comparison of all multi-tissue methods. *We applied SWAM to all combinations of GTEx and DGN resources. For the GTEx resources, we always used every tissue available. In version 6, this comprised of 44 tissues. For version 7, there were 48 tissues and version 8 contained 49 tissues. For the sake of consistency, our target tissue for each of these combinations was GTEx v6 LCL.*

| Method | Total #<br>genes | Genes with<br>FDR < 0.05 | P-value<br>threshold for<br>FDR=0.05 | Genes with<br>p-value < 0.05 |
| --- | --- | --- | --- | --- |
| SWAM-LCL (GTEx v6) | 13213 | 3040 | 0.01145 | 4148 |
| SWAM-LCL (GTEx v6 + DGN) | 13213 | 3192 | 0.012077 | 4301 |
| SWAM-LCL (GTEx v7) | 13213 | 3060 | 0.011463 | 4215 |
| SWAM-LCL (GTEx v7 + DGN) | 13213 | 3411 | 0.012903 | 4469 |
| SWAM-LCL (GTEx v8) | 13213 | 3203 | 0.01212 | 4236 |
| SWAM-LCL (GTEx v8 + DGN) | 13213 | 3449 | 0.013046 | 4460 |
| SWAM-LCL (GTEx v6 + v7) | 13213 | 3283 | 0.012383 | 4385 |
| SWAM-LCL (GTEx v6 + v7 + DGN) | 13213 | 3361 | 0.012674 | 4480 |
| SWAM-LCL (GTEx v6 + v8) | 13213 | 3134 | 0.01185 | 4274 |
| SWAM-LCL (GTEx v6 + v8 + DGN) | 13213 | 3275 | 0.012389 | 4384 |
| SWAM-LCL (GTEx v7 + v8) | 13213 | 3259 | 0.01227 | 4326 |
| SWAM-LCL (GTEx v7 + v8 + DGN) | 13213 | 3368 | 0.012737 | 4478 |
| SWAM-LCL (GTEx v6 + v7 + v8) | 13213 | 3342 | 0.012641 | 4448 |
| SWAM-LCL (GTEx v6 + v7 + v8 +<br>DGN) | 13213 | 3413 | 0.012878 | 4526 |
| UTMOST-LCL | 13213 | 2238 | 0.008466 | 3185 |
| NAIVE AVERAGE | 13213 | 2666 | 0.010066 | 3830 |
| BEST TISSUE | 13213 | 2493 | 0.009394 | 3663 |

**Supplementary Table 9 –.** comparison of GTEx v7/v8 single tissue models versus GEUVADIS LCL. *We also compared every prediXcan model derived from GTEx version 7 and version 8 tissues, and tested imputation accuracy against GEUVADIS LCL measured expression levels. Surprisingly, despite the increase in sample size, the LCL tissue from v8 performed worse than its version 7 counterpart. The number of tissues outperforming LCL in both v7 and v8 highlight the opportunity to leverage information from other tissues to improve imputation accuracy for under-powered tissues.*

| Tissue | GTEx v7 |  |  | GTEx v8 |  |  |
| --- | --- | --- | --- | --- | --- | --- |
|  | sample size | Total # genes | Genes with FDR < 0.05 | sample size | Total # genes | Genes with FDR < 0.05 |
| Adipose Subcutaneous | 328 | 6689 | 2259 | 581 | 6847 | 2114 |
| Adipose Visceral Omentum | 273 | 5286 | 1905 | 469 | 5769 | 1957 |
| Adrenal Gland | 146 | 3784 | 1322 | 233 | 3816 | 1279 |
| Artery Aorta | 236 | 5488 | 1850 | 387 | 6090 | 1844 |
| Artery Coronary | 128 | 2838 | 1040 | 213 | 3177 | 1119 |
| Artery Tibial | 329 | 6836 | 2131 | 584 | 6955 | 2023 |
| Brain Amygdala | 81 | 1876 | 581 | 129 | 2081 | 682 |
| Brain Anterior cingulate cortex BA24 | 102 | 2628 | 838 | 147 | 2662 | 840 |
| Brain Caudate basal ganglia | 126 | 3272 | 1035 | 194 | 3795 | 1146 |
| Brain Cerebellar Hemisphere | 113 | 3810 | 1050 | 175 | 4482 | 1127 |
| Brain Cerebellum | 137 | 4899 | 1310 | 209 | 5254 | 1299 |
| Brain Cortex | 119 | 3422 | 1073 | 205 | 4169 | 1203 |
| Brain Frontal Cortex BA9 | 104 | 2812 | 879 | 175 | 3424 | 1019 |
| Brain Hippocampus | 99 | 2217 | 708 | 165 | 2806 | 886 |
| Brain Hypothalamus | 98 | 2219 | 710 | 170 | 2742 | 911 |
| Brain Nucleus accumbens basal ganglia | 114 | 2820 | 921 | 202 | 3629 | 1076 |
| Brain Putamen basal ganglia | 98 | 2542 | 806 | 170 | 3365 | 1035 |
| Brain Spinal cord cervical c-1 | 76 | 2003 | 600 | 126 | 2455 | 744 |
| Brain Substantia nigra | 70 | 1609 | 496 | 114 | 1892 | 570 |
| Breast Mammary Tissue | 211 | 4280 | 1622 | 396 | 5076 | 1756 |
| Cells EBV-transformed lymphocytes | 96 | 2777 | 1731 | 147 | 2537 | 1620 |
| Cells Transformed fibroblasts | 256 | 6297 | 2336 | 483 | 7421 | 2428 |
| Colon Sigmoid | 185 | 4257 | 1556 | 318 | 4847 | 1633 |
| Colon Transverse | 210 | 4457 | 1771 | 368 | 4923 | 1781 |
| Esophagus Gastroesophageal Junction | 185 | 4325 | 1603 | 330 | 4964 | 1675 |
| Esophagus Mucosa | 307 | 6744 | 2305 | 497 | 6872 | 2167 |
| Esophagus Muscularis | 287 | 6354 | 2119 | 465 | 6554 | 2030 |
| Heart Atrial Appendage | 231 | 4866 | 1675 | 372 | 5262 | 1696 |
| Heart Left Ventricle | 233 | 4449 | 1491 | 386 | 4902 | 1569 |
| Kidney Cortex | NA | NA | NA | 73 | 1205 | 344 |
| Liver | 134 | 2746 | 883 | 208 | 2983 | 976 |
| Lung | 333 | 6251 | 2190 | 515 | 6173 | 2071 |
| Minor Salivary Gland | 74 | 1785 | 652 | 144 | 2161 | 842 |
| Muscle Skeletal | 421 | 6323 | 1901 | 706 | 6261 | 1762 |
| Nerve Tibial | 305 | 7512 | 2179 | 532 | 7764 | 2051 |
| Ovary | 99 | 2406 | 818 | 167 | 2751 | 909 |
| Pancreas | 180 | 4369 | 1504 | 305 | 4710 | 1532 |
| Pituitary | 143 | 3691 | 1228 | 237 | 4262 | 1381 |
| Prostate | 114 | 2487 | 926 | 221 | 3205 | 1125 |
| Skin Not Sun Exposed Suprapubic | 285 | 6092 | 1997 | 517 | 6802 | 2011 |
| Skin Sun Exposed Lower leg | 359 | 7221 | 2201 | 605 | 7203 | 2071 |
| Small Intestine Terminal Ileum | 103 | 2482 | 991 | 174 | 2844 | 1145 |
| Spleen | 119 | 3753 | 1514 | 227 | 4527 | 1720 |
| Stomach | 200 | 3899 | 1518 | 324 | 4064 | 1542 |
| Testis | 191 | 5919 | 1451 | 322 | 6470 | 1518 |
| Thyroid | 344 | 7556 | 2249 | 574 | 7468 | 2130 |
| Uterus | 82 | 1957 | 681 | 129 | 1944 | 699 |
| Vagina | 91 | 1889 | 690 | 141 | 1919 | 725 |
| Whole Blood | 315 | 5432 | 1915 | 670 | 6195 | 2082 |

**Supplementary Table 10 –.**
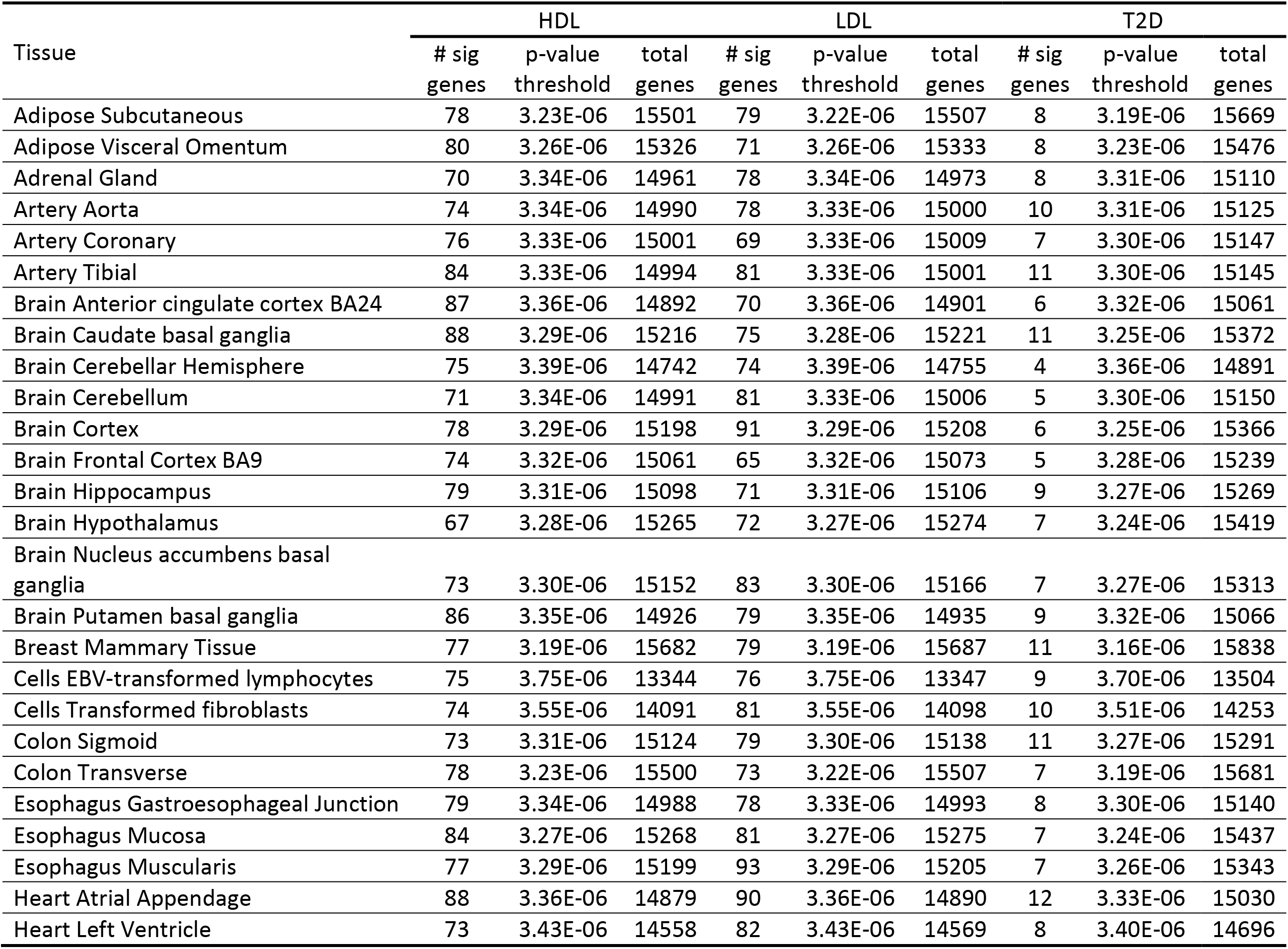
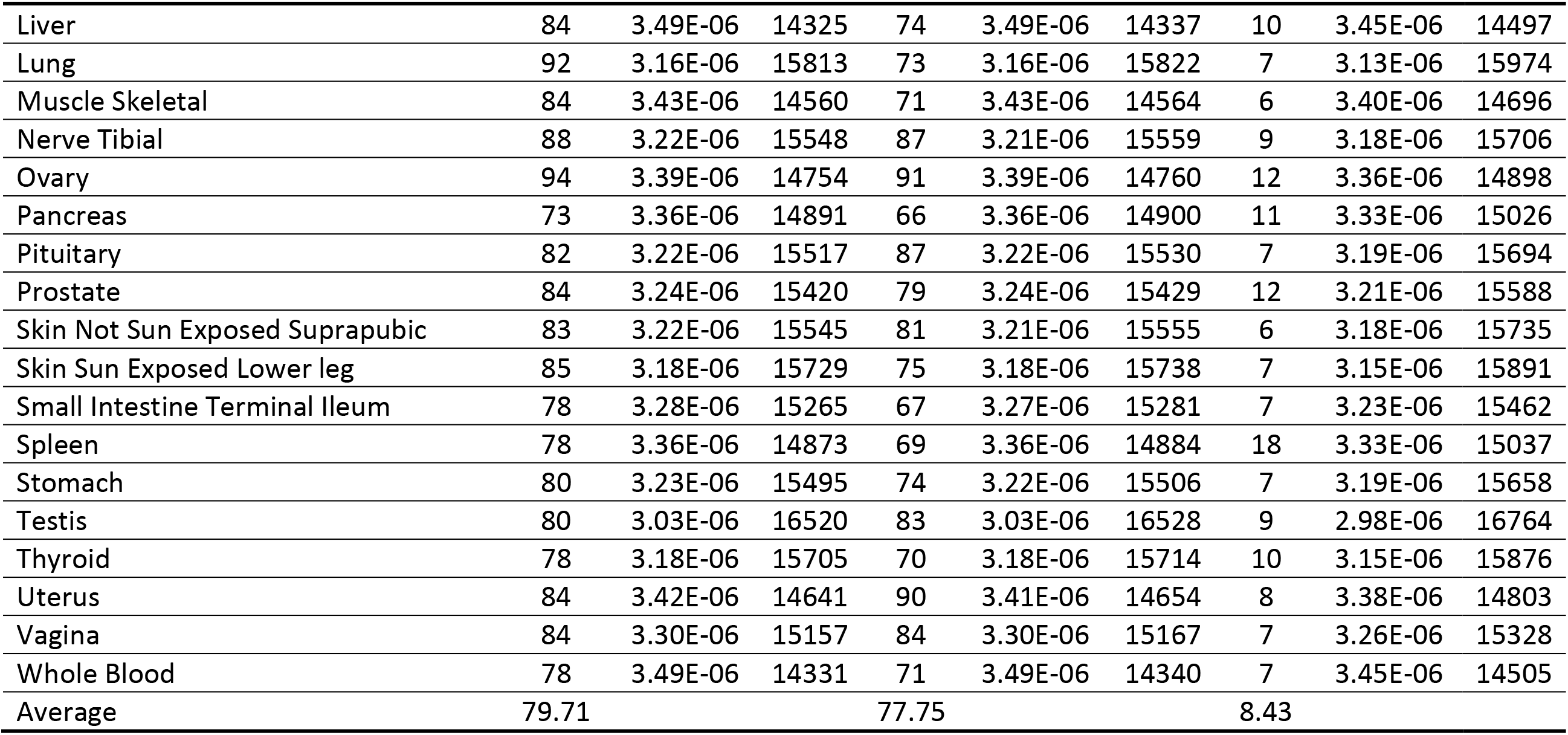
TWAS association signals for SWAM. *We used SWAM to derive an tissue-specific model for every GTEx version 6 tissue, and used these models as inputs to metaXcan to infer TWAS results. As mentioned in the methods section, the HDL and LDL traits were from Global Lipids Genetics Consortium (GLGC) and Type-2 Diabetes (T2D) from the DIAGRAM consortium.*

**Supplementary Table 11 –.** TWAS association signals for UTMOST. *These models were also derived from GTEx version 6 tissues using the UTMOST method. Models were downloaded from https://github.com/Joker-Jerome/UTMOST*

| Tissue | HDL |  |  | LDL |  |  | T2D |  |  |
| --- | --- | --- | --- | --- | --- | --- | --- | --- | --- |
|  | # sig genes | p-value threshold | total genes | # sig genes | p-value threshold | total genes | # sig genes | p-value threshold | total genes |
| Adipose Subcutaneous | 82 | 4.18E-06 | 11964 | 61 | 4.18E-06 | 11969 | 11 | 3.94E-06 | 12688 |
| Adipose Visceral Omentum | 69 | 4.26E-06 | 11741 | 68 | 4.26E-06 | 11743 | 10 | 4.01E-06 | 12475 |
| Adrenal Gland | 72 | 4.55E-06 | 10998 | 66 | 4.54E-06 | 11004 | 7 | 4.26E-06 | 11737 |
| Artery Aorta | 75 | 4.47E-06 | 11180 | 55 | 4.47E-06 | 11184 | 7 | 4.19E-06 | 11926 |
| Artery Coronary | 61 | 4.39E-06 | 11391 | 58 | 4.39E-06 | 11397 | 7 | 4.13E-06 | 12114 |
| Artery Tibial | 74 | 4.43E-06 | 11292 | 60 | 4.43E-06 | 11295 | 9 | 4.15E-06 | 12044 |
| Brain Anterior cingulate cortex BA24 | 53 | 5.00E-06 | 10002 | 62 | 5.00E-06 | 10003 | 8 | 4.64E-06 | 10779 |
| Brain Caudate basal ganglia | 61 | 4.61E-06 | 10850 | 55 | 4.61E-06 | 10850 | 11 | 4.30E-06 | 11627 |
| Brain Cerebellar Hemisphere | 65 | 4.94E-06 | 10113 | 64 | 4.94E-06 | 10115 | 11 | 4.60E-06 | 10864 |
| Brain Cerebellum | 62 | 4.78E-06 | 10457 | 67 | 4.78E-06 | 10457 | 7 | 4.46E-06 | 11205 |
| Brain Cortex | 61 | 4.69E-06 | 10650 | 57 | 4.70E-06 | 10647 | 8 | 4.37E-06 | 11445 |
| Brain Frontal Cortex BA9 | 66 | 4.67E-06 | 10717 | 57 | 4.66E-06 | 10719 | 11 | 4.34E-06 | 11519 |
| Brain Hippocampus | 58 | 4.67E-06 | 10701 | 54 | 4.67E-06 | 10701 | 8 | 4.36E-06 | 11469 |
| Brain Hypothalamus | 75 | 4.55E-06 | 10986 | 65 | 4.55E-06 | 10987 | 11 | 4.25E-06 | 11764 |
| Brain Nucleus accumbens basal ganglia | 74 | 4.64E-06 | 10781 | 59 | 4.64E-06 | 10783 | 9 | 4.33E-06 | 11541 |
| Brain Putamen basal ganglia | 71 | 4.84E-06 | 10323 | 55 | 4.84E-06 | 10327 | 11 | 4.49E-06 | 11129 |
| Breast Mammary Tissue | 74 | 4.12E-06 | 12141 | 62 | 4.12E-06 | 12143 | 8 | 3.88E-06 | 12899 |
| Cells EBV-transformed lymphocytes | 65 | 5.20E-06 | 9610 | 50 | 5.20E-06 | 9615 | 5 | 4.87E-06 | 10267 |
| Cells Transformed fibroblasts | 61 | 4.80E-06 | 10406 | 60 | 4.80E-06 | 10409 | 10 | 4.51E-06 | 11089 |
| Colon Sigmoid | 72 | 4.44E-06 | 11257 | 68 | 4.44E-06 | 11261 | 7 | 4.16E-06 | 12010 |
| Colon Transverse | 74 | 4.25E-06 | 11759 | 61 | 4.25E-06 | 11765 | 10 | 3.99E-06 | 12545 |
| Esophagus Gastroesophageal Junction | 58 | 4.50E-06 | 11117 | 52 | 4.50E-06 | 11119 | 8 | 4.23E-06 | 11823 |
| Esophagus Mucosa | 77 | 4.31E-06 | 11595 | 71 | 4.31E-06 | 11603 | 7 | 4.06E-06 | 12316 |
| Esophagus Muscularis | 67 | 4.43E-06 | 11285 | 67 | 4.43E-06 | 11289 | 7 | 4.16E-06 | 12029 |
| Heart Atrial Appendage | 69 | 4.58E-06 | 10922 | 64 | 4.58E-06 | 10923 | 9 | 4.28E-06 | 11679 |
| Heart Left Ventricle | 77 | 4.75E-06 | 10519 | 55 | 4.75E-06 | 10522 | 10 | 4.45E-06 | 11242 |
| Liver | 63 | 4.97E-06 | 10062 | 69 | 4.97E-06 | 10068 | 7 | 4.63E-06 | 10808 |
| Lung | 81 | 4.02E-06 | 12428 | 62 | 4.02E-06 | 12433 | 8 | 3.80E-06 | 13152 |
| Muscle Skeletal | 81 | 4.70E-06 | 10629 | 65 | 4.70E-06 | 10631 | 6 | 4.41E-06 | 11326 |
| Nerve Tibial | 78 | 4.24E-06 | 11784 | 63 | 4.24E-06 | 11787 | 7 | 4.00E-06 | 12511 |
| Ovary | 57 | 4.71E-06 | 10608 | 60 | 4.71E-06 | 10607 | 8 | 4.42E-06 | 11323 |
| Pancreas | 60 | 4.71E-06 | 10619 | 63 | 4.71E-06 | 10625 | 7 | 4.38E-06 | 11410 |
| Pituitary | 75 | 4.41E-06 | 11336 | 69 | 4.41E-06 | 11337 | 8 | 4.13E-06 | 12117 |
| Prostate | 73 | 4.28E-06 | 11685 | 60 | 4.28E-06 | 11688 | 8 | 4.02E-06 | 12450 |
| Skin Not Sun Exposed Suprapubic | 75 | 4.24E-06 | 11789 | 69 | 4.24E-06 | 11797 | 7 | 4.00E-06 | 12505 |
| Skin Sun Exposed Lower leg | 72 | 4.12E-06 | 12144 | 62 | 4.12E-06 | 12149 | 8 | 3.87E-06 | 12904 |
| Small Intestine Terminal Ileum | 68 | 4.48E-06 | 11150 | 64 | 4.48E-06 | 11151 | 10 | 4.19E-06 | 11938 |
| Spleen | 70 | 4.66E-06 | 10736 | 55 | 4.66E-06 | 10739 | 13 | 4.36E-06 | 11474 |
| Stomach | 78 | 4.23E-06 | 11833 | 68 | 4.22E-06 | 11838 | 14 | 3.97E-06 | 12580 |
| Testis | 73 | 4.03E-06 | 12411 | 70 | 4.03E-06 | 12412 | 7 | 3.78E-06 | 13222 |
| Thyroid | 82 | 4.13E-06 | 12106 | 68 | 4.13E-06 | 12113 | 17 | 3.88E-06 | 12873 |
| Uterus | 60 | 4.93E-06 | 10148 | 61 | 4.93E-06 | 10151 | 8 | 4.62E-06 | 10813 |
| Vagina | 70 | 4.43E-06 | 11285 | 59 | 4.43E-06 | 11288 | 7 | 4.16E-06 | 12018 |
| Whole Blood | 59 | 4.76E-06 | 10511 | 52 | 4.75E-06 | 10518 | 12 | 4.48E-06 | 11168 |
| Average | 69.27 |  |  | 61.64 |  |  | 8.84 |  |  |

**Supplementary Table 12 –.** TWAS association signals for prediXcan (single-tissue) *TWAS results via metaXcan using prediXcan single tissue models derived from GTEx version 6 tissues*

| Tissue | HDL |  |  | LDL |  |  | T2D |  |  |
| --- | --- | --- | --- | --- | --- | --- | --- | --- | --- |
|  | # sig genes | p-value threshold | total genes | # sig genes | p-value threshold | total genes | # sig genes | p-value threshold | total genes |
| TW Adipose Subcutaneous | 46 | 7.54E-06 | 6634 | 28 | 7.54E-06 | 6628 | 10 | 6.91E-06 | 7234 |
| TW Adipose Visceral Omentum | 32 | 1.20E-05 | 4174 | 21 | 1.20E-05 | 4173 | 6 | 1.10E-05 | 4542 |
| TW Adrenal Gland | 23 | 1.33E-05 | 3760 | 20 | 1.33E-05 | 3758 | 3 | 1.24E-05 | 4048 |
| TW Artery Aorta | 33 | 8.76E-06 | 5706 | 44 | 8.76E-06 | 5705 | 5 | 8.11E-06 | 6163 |
| TW Artery Coronary | 13 | 1.65E-05 | 3025 | 21 | 1.65E-05 | 3024 | 3 | 1.54E-05 | 3242 |
| TW Artery Tibial | 36 | 7.50E-06 | 6666 | 32 | 7.51E-06 | 6657 | 6 | 6.89E-06 | 7259 |
| TW Brain Anterior cingulate cortex BA24 | 6 | 2.03E-05 | 2466 | 14 | 2.03E-05 | 2466 | 2 | 1.88E-05 | 2654 |
| TW Brain Caudate basal ganglia | 21 | 1.48E-05 | 3375 | 25 | 1.48E-05 | 3372 | 2 | 1.38E-05 | 3616 |
| TW Brain Cerebellar Hemisphere | 12 | 1.26E-05 | 3955 | 18 | 1.26E-05 | 3954 | 3 | 1.18E-05 | 4228 |
| TW Brain Cerebellum | 22 | 1.10E-05 | 4543 | 28 | 1.10E-05 | 4542 | 4 | 1.04E-05 | 4830 |
| TW Brain Cortex | 19 | 1.49E-05 | 3351 | 33 | 1.49E-05 | 3347 | 3 | 1.40E-05 | 3583 |
| TW Brain Frontal Cortex BA9 | 12 | 1.66E-05 | 3013 | 16 | 1.66E-05 | 3013 | 1 | 1.56E-05 | 3211 |
| TW Brain Hippocampus | 12 | 2.12E-05 | 2362 | 10 | 2.12E-05 | 2362 | 1 | 1.97E-05 | 2534 |
| TW Brain Hypothalamus | 6 | 2.20E-05 | 2269 | 11 | 2.20E-05 | 2268 | 1 | 2.04E-05 | 2455 |
| TW Brain Nucleus accumbens basal ganglia | 16 | 1.70E-05 | 2935 | 17 | 1.70E-05 | 2935 | 1 | 1.60E-05 | 3131 |
| TW Brain Putamen basal ganglia | 14 | 1.91E-05 | 2620 | 17 | 1.91E-05 | 2620 | 5 | 1.78E-05 | 2807 |
| TW Breast Mammary Tissue | 23 | 1.16E-05 | 4292 | 20 | 1.17E-05 | 4288 | 4 | 1.07E-05 | 4655 |
| TW Cells EBV-transformed lymphocytes | 22 | 1.43E-05 | 3493 | 17 | 1.43E-05 | 3491 | 3 | 1.33E-05 | 3751 |
| TW Cells Transformed fibroblasts | 52 | 7.02E-06 | 7120 | 35 | 7.03E-06 | 7114 | 7 | 6.50E-06 | 7692 |
| TW Colon Sigmoid | 16 | 1.40E-05 | 3561 | 16 | 1.40E-05 | 3559 | 4 | 1.30E-05 | 3858 |
| TW Colon Transverse | 24 | 1.11E-05 | 4485 | 24 | 1.12E-05 | 4480 | 4 | 1.03E-05 | 4874 |
| TW Esophagus Gastroesophageal Junction | 14 | 1.45E-05 | 3456 | 17 | 1.45E-05 | 3454 | 3 | 1.34E-05 | 3727 |
| TW Esophagus Mucosa | 38 | 7.78E-06 | 6425 | 32 | 7.78E-06 | 6423 | 3 | 7.18E-06 | 6961 |
| TW Esophagus Muscularis | 31 | 8.37E-06 | 5971 | 33 | 8.38E-06 | 5966 | 4 | 7.70E-06 | 6493 |
| TW Heart Atrial Appendage | 28 | 1.19E-05 | 4187 | 21 | 1.19E-05 | 4185 | 6 | 1.11E-05 | 4501 |
| TW Heart Left Ventricle | 25 | 1.11E-05 | 4517 | 33 | 1.11E-05 | 4513 | 4 | 1.02E-05 | 4885 |
| TW Liver | 16 | 1.86E-05 | 2695 | 19 | 1.86E-05 | 2692 | 4 | 1.72E-05 | 2909 |
| TW Lung | 42 | 8.32E-06 | 6008 | 16 | 8.33E-06 | 6002 | 5 | 7.56E-06 | 6611 |
| TW Muscle Skeletal | 30 | 8.32E-06 | 6009 | 30 | 8.33E-06 | 6003 | 8 | 7.56E-06 | 6614 |
| TW Nerve Tibial | 41 | 6.60E-06 | 7577 | 39 | 6.61E-06 | 7570 | 6 | 6.13E-06 | 8157 |
| TW Ovary | 13 | 1.94E-05 | 2576 | 17 | 1.94E-05 | 2575 | 3 | 1.81E-05 | 2762 |
| TW Pancreas | 26 | 1.12E-05 | 4449 | 26 | 1.12E-05 | 4447 | 4 | 1.05E-05 | 4771 |
| TW Pituitary | 12 | 1.59E-05 | 3145 | 13 | 1.59E-05 | 3144 | 5 | 1.48E-05 | 3382 |
| TW Prostate | 10 | 2.09E-05 | 2389 | 20 | 2.09E-05 | 2389 | 3 | 1.92E-05 | 2609 |
| TW Skin Not Sun Exposed Suprapubic | 27 | 9.37E-06 | 5336 | 31 | 9.38E-06 | 5333 | 4 | 8.62E-06 | 5798 |
| TW Skin Sun Exposed Lower leg | 42 | 7.10E-06 | 7041 | 35 | 7.11E-06 | 7037 | 6 | 6.55E-06 | 7628 |
| TW Small Intestine Terminal Ileum | 15 | 1.97E-05 | 2538 | 16 | 1.97E-05 | 2536 | 3 | 1.83E-05 | 2729 |
| TW Spleen | 19 | 1.45E-05 | 3456 | 18 | 1.45E-05 | 3456 | 5 | 1.35E-05 | 3698 |
| TW Stomach | 16 | 1.27E-05 | 3945 | 23 | 1.27E-05 | 3943 | 4 | 1.17E-05 | 4271 |
| TW Testis | 30 | 7.37E-06 | 6780 | 28 | 7.38E-06 | 6778 | 7 | 6.91E-06 | 7234 |
| TW Thyroid | 39 | 6.69E-06 | 7478 | 40 | 6.69E-06 | 7469 | 7 | 6.14E-06 | 8140 |
| TW Uterus | 11 | 2.51E-05 | 1991 | 8 | 2.51E-05 | 1991 | 1 | 2.34E-05 | 2139 |
| TW Vagina | 15 | 2.57E-05 | 1946 | 10 | 2.57E-05 | 1944 | 1 | 2.40E-05 | 2082 |
| TW Whole Blood | 42 | 8.23E-06 | 6077 | 30 | 8.23E-06 | 6073 | 3 | 7.42E-06 | 6743 |
| Average | 23.68 |  |  | 23.23 |  |  | 4.02 |  |  |

